# Exponential Map Models as an Interpretable Framework for Generating Neural Spatial Representations

**DOI:** 10.1101/2025.09.25.678504

**Authors:** Markus Pettersen, Nicolai Haug, Joakim Bergli, Thomas M. Surowiec, Mikkel Lepperød

## Abstract

A foundational challenge in neuroscience and AI is understanding how physical space is mapped into neural representations. While artificial neural networks can learn brain-like spatial representations, their “black-box” nature obscures whether these representations reflect general principles or artifacts of specific architectures, objectives or optimization choices, offering no guarantee that solutions for tasks like path integration generalize beyond training data. Here, we use a first-principles framework based on the matrix exponential, in which generator matrices map physical locations to neural population vectors without gradient-based optimization. Within this formalism, navigational properties follow directly from algebraic constraints on the generators: commuting generators enable trajectory-invariant, exact path integration, while skew-symmetry yields equinorm representations with translation-invariant similarities. Preserving the flat Euclidean metric further constrains the generator spectra to balanced wavevector configurations, such as the roots-of-unity constellations that generate periodic grid fields. Similarity preservation further produces localized place-like fields, and extending the generators to a context signal models context-dependent remapping. Beyond its algebraic relation to continuous attractor and oscillatory interference models, the framework natively supports goal-oriented multi-map navigation through hyperdimensional bundling and binding, fits recorded grid and place cell data, extrapolates their rate maps far beyond the fitted region, and makes testable predictions for entorhinal circuits. By grounding spatial representations in exact algebraic conditions, this work offers a transparent alternative to empirical representation learning, revealing the conditions under which a coherent neural map of space arises.

## 1 Introduction

A fundamental problem in neuroscience and artificial intelligence (AI) is understanding the mapping from physical space to the representational space of neural population activity. In the mammalian brain, this mapping is supported by the hippocampal-entorhinal system. This circuit contains place cells [1], which fire in specific localized areas, and grid cells [2], whose striking periodic hexagonal firing patterns are hypothesized to provide a neural metric for space [3, 4, 5]. Together with other spatially tuned neurons, such as head direction [6] and border cells [7, 8, 9], they form a high-dimensional representation of an animal’s location and heading. This spatial map can abruptly reorganize in response to contextual changes, a phenomenon known as remapping [10, 11], demonstrating that neurons also encode environmental identity.

To explain the hexagonal firing patterns of grid cells, theoretical neuroscience historically relied on hand-crafted mechanistic models. Continuous attractor neural networks (CANNs) posit that grid fields arise from a periodic activity pattern shaped by recurrent inhibition across a neural sheet, which velocity inputs translate as the animal moves [12]. Conversely, oscillatory interference (OI) models propose that spatial periodicity emerges from thresholded interference between temporal velocity-controlled oscillators within individual neurons [13, 14]. While these models make their mechanisms explicit, the geometry of the representation is built into the recurrent connectivity or the oscillator tuning by hand. As a result, they cannot show that grid-like representations arise as a solution to navigation, nor can they readily learn such representations from data. More recently, deep recurrent neural networks (RNNs) trained to solve navigation tasks have empirically learned representations resembling biological place and grid cells [15, 16, 17, 18, 19]. Self-supervised learning architectures have demonstrated that periodic spatial tuning is a normative solution to specific geometric constraints [20, 21]. However, the black-box nature of deep neural networks makes it difficult to disentangle whether learned representations reflect fundamental principles of navigation or are artifacts of a chosen architecture, objective function, or training protocol. Consequently, data-driven methodologies offer no algebraic guarantee that navigational solutions will generalize beyond their training environments.

In contrast, recent theoretical work has begun to derive properties of spatial representations by framing displacements in space as group actions on the neural population vector [22, 23, 24]. Within this formulation, conformal isometry models [22, 23, 25] show that hexagonal grid patterns emerge when distance preservation is imposed, but obtain their representations through gradient-based training. Written as matrices, these actions take the form of a matrix exponential, whose generator matrices are learned together with the representation [22]. Work on actionable representations [24] shows that such a group action decomposes the representation into two-dimensional rotational subspaces, while the frequencies of these subspaces are found by numerical optimization. Fourier models of grid cells [26, 27, 28] and spatial semantic pointers [29, 30] similarly represent position using phase-shifted sinusoids, with wavevectors imposed by hand or drawn at random. In all of these models, the wavevectors that shape the representation are learned or chosen rather than derived, and the conditions they must satisfy for the resulting map of space to be coherent have not been given in closed form.

Here, we provide these conditions within a first-principles framework based on the matrix exponential, in which generator matrices map physical locations to neural population vectors (Fig. 1a,b) and the representations are constructed analytically rather than trained. Although the resulting representations are superpositions of plane waves, operating on the generators directly allows us to pose path invariance, translational invariance, and metric preservation as algebraic constraints and solve them exactly. By the Baker-Campbell-Hausdorff formula, path integration is exact when the generators commute (Fig. 1c), and when they do not, the leading error grows with the commutator and with the area enclosed by the path. This recovers the commutativity condition established for continuous linear systems [31] within the matrix exponential setting. Skew-symmetric generators give equinorm representations, and we show that their similarities are then translationally invariant (Fig. 1d). Furthermore, we derive a closed-form condition under which the representation preserves the flat Euclidean metric (Fig. 1e). The spatial wavevectors set up by the generator eigenvalues must form a balanced configuration. In such a configuration the wavevectors are spread over orientations such that the representation stretches space by the same factor in every direction. Sets of roots of unity on discrete rings in frequency space are balanced, and with only two or three wavevectors of equal amplitude they are the only balanced configurations. The set with three wavevectors produces hexagonal grid fields. The generators also connect our work to CANNs and OI models, as these matrices map onto velocity-modulated recurrent connectivity while their spectra describe interfering oscillators. When the geometric conditions are met, the framework bridges the two analytically, with periodic grid fields emerging as a solution rather than being assumed. Finally, since the algebra extends naturally to non-spatial variables, adding a context generator produces context-dependent remapping within the same construction (Fig. 1f), and the model supports reward-based multi-map navigation using principles from hyperdimensional computing [32].

**Figure 1:**
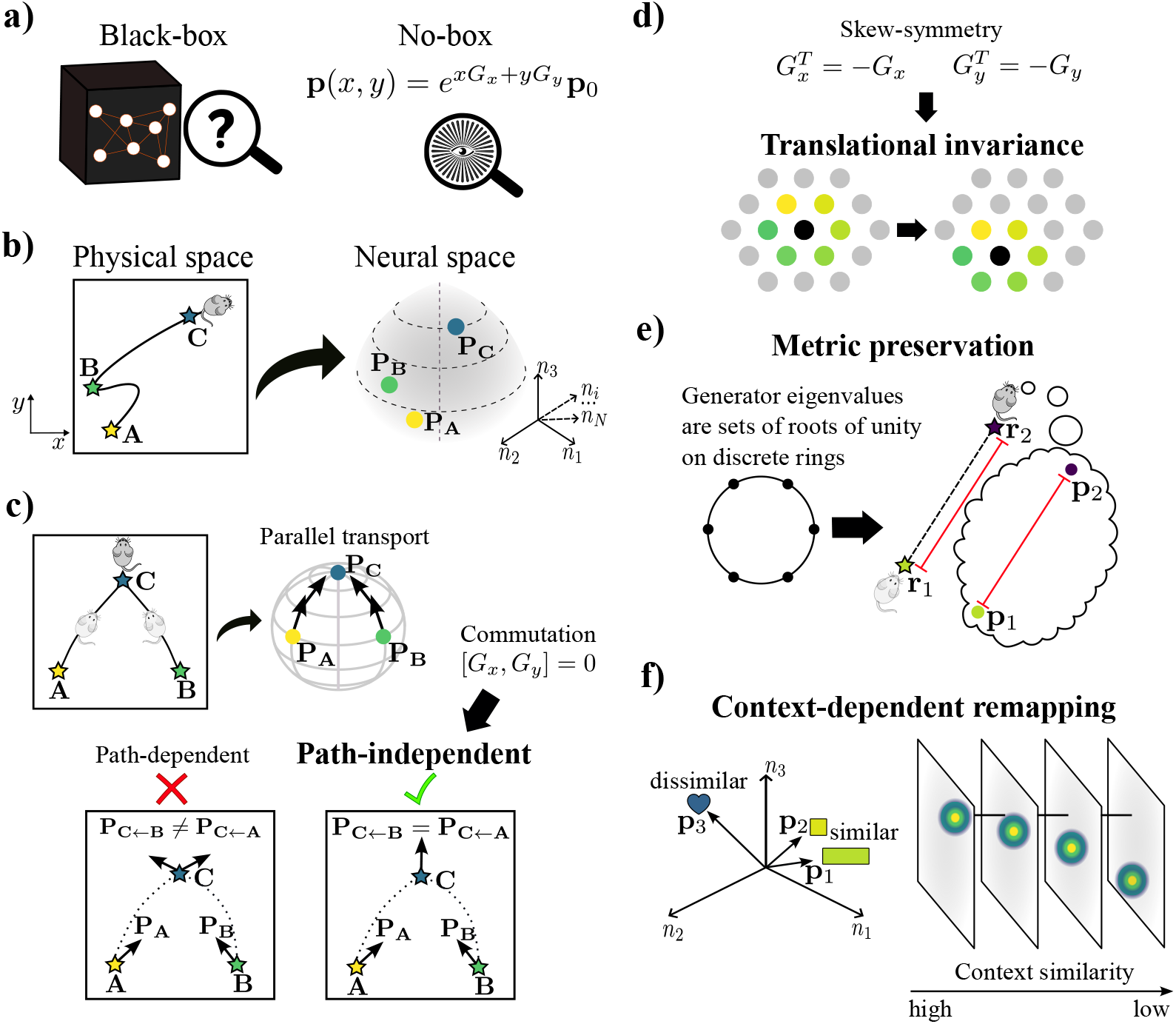
The exponential map framework for interpretable spatial representations. **a)** The exponential map model is a transparent “no-box” alternative to deep learning models, using generator matrices (*G*_*x*_, *G*_*y*_) to construct a neural population vector **p**(*x, y*). **b)** A population vector is assigned to every location, mapping physical space to neural space. **c) Path invariance:** Exact path integration requires representations to be independent of the traversal route. A path-dependent traversal yields unequal representations (**p**_*C*_ *←*_*B*_ ≠ **p**_*C← A*_, red cross), whereas path invariance is guaranteed if the generators commute ([*G*_*x*_, *G*_*y*_] = 0, green checkmark). **d) Translational invariance:** Skew-symmetric generators (*G*^*T*^ = *G*) yield orthogonal transformations, maintaining a constant vector norm (equinorm constraint) and ensuring representational similarities depend solely on displacement. **e)Metric preservation:** The flat metric of physical space is preserved in neural space when the generator eigenvalues form balanced configurations, such as roots of unity on discrete rings in frequency space. **f) Remapping:** A context generator extends the framework to non-spatial variables. Representations of similar contexts remain similar, while those of dissimilar contexts decorrelate.

## 2 Results and discussion

### 2.1 Constructing spatial representations

A spatial representation is a mapping that assigns a neural population vector **p**(*x, y*) ∈ ℝ^*N*^ to a physical coordinate (*x, y*). Each of the *N* components represents a neuron’s firing rate,^1^ so that **p** is a point in an *N* -dimensional state space. Building upon previous models [22, 33], we construct the representation with the matrix exponential:

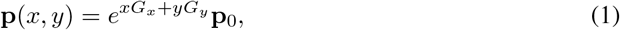

where *G*_*x*_, *G*_*y*_ ∈ ℝ^*N* ×*N*^ are generator matrices for the two cardinal directions, and **p**_0_ is the population vector at the origin. The generator matrices define how locations in physical space are translated into transformations in the high-dimensional neural state space. The matrix exponential composes the transformations to transport **p**_0_ to any target location (*x, y*). Arbitrary generators, however, can cause the representation’s norm to collapse or diverge, or make the state reached by path integration path-dependent. Because this framework is mathematically transparent, we can derive exact algebraic conditions on *G*_*x*_ and *G*_*y*_ that guarantee properties essential to navigation.

### 2.2 Path integration and path independence

A spatial map must support the computations navigation relies on. One of the most fundamental is path integration, in which a navigator estimates its position by integrating self-motion cues. Path integration requires that the representation at a new location follows from the current state and a displacement operator *Q*(Δ*x*, Δ*y*) alone, such that **p**(*x* + Δ*x, y* + Δ*y*) = *Q*(Δ*x*, Δ*y*)**p**(*x, y*). For a displacement from the origin, Eq. (1) shows that this holds with the operator

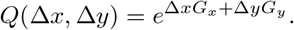

From any other location, it holds only if applying two displacements in succession equals applying their sum. By the Baker-Campbell-Hausdorff expansion, this is the case exactly when the generators commute, [*G*_*x*_, *G*_*y*_] = 0 (see Appendix B). This is the discrete counterpart of the result that exact path integration in continuous linear systems requires commuting transition matrices [31]. With commuting generators, path integration is therefore exact and independent of the route taken, and velocity need not be constant.

In biological circuits, exact commutativity is unlikely to hold, since synaptic noise and anisotropic connectivity could perturb effective generators. For non-commuting generators, the Baker-Campbell-Hausdorff expansion acquires a leading correction proportional to (Δ*x*_1_Δ*y*_2_ − Δ*y*_1_Δ*x*_2_)[*G*_*x*_, *G*_*y*_] for consecutive displacements (Δ*x*_1_, Δ*y*_1_) and (Δ*x*_2_, Δ*y*_2_), that is, to the commutator times the area spanned by the two path segments (see Appendix B). The commutator norm [∥*G*_*x*_, *G*_*y*_∥] would thus provide an analytical measure of trajectory-dependent drift, and sensory anchoring to boundaries or place-like inputs may be required to reset the state to a reference. However, this characterization is analytical, and robustness under realistic navigation conditions remains an open question. In what follows we assume commutativity and show that it allows the similarity structure of the representation to be constructed explicitly.

### 2.3 Orthogonal transformations enable egocentric navigation

To navigate the open field, an animal must be able to disambiguate neural population vectors of distinct locations. We therefore ask how the similarity between two representations depends on where they are in the environment. As a measure, we use the cosine similarity (normalized inner product) between a state at (*x, y*) and a displaced state at (*x* + Δ*x, y* + Δ*y*),

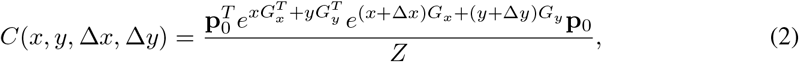

where *Z* is the normalization factor.

If the generator matrices *G*_*x*_ and *G*_*y*_ are skew-symmetric (*G*^*T*^ = −*G*), two properties follow. First, the matrix exponential of a skew-symmetric matrix is orthogonal. The transformation therefore preserves the norm of the population vector everywhere (∥**p**(*x, y*) ∥ _2_ = ∥**p**_0_∥ _2_), which fixes the normalization factor to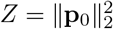. Second, with *G*^*T*^ = − *G* the first exponential in Eq. (2) equals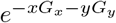, and since the generators commute, as assumed for path integration in Section 2.2, the two exponentials combine into one and the position-dependent terms cancel. The similarity is then translation invariant, depending only on the displacement (Δ*x*, Δ*y*),

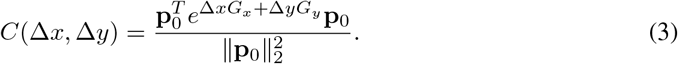

Translational invariance enables relative or egocentric navigation, as absolute position is not needed to judge the similarity between representations. Skew-symmetric generators may therefore be useful for open-field navigation, where an animal’s location may be unknown. CANNs obtain the same property through translation-invariant connectivity, which lets the activity pattern shift rigidly as the animal moves [12, 3]. Several trained models impose an equinorm constraint through a regularizer or an architectural constraint [20, 23, 19], whereas here it follows from skew-symmetry. The same condition also gives translational invariance, which suggests why such a constraint is computationally useful.

By the spectral theorem for real skew-symmetric matrices, the commuting generators can be brought simultaneously into canonical block diagonal form, *G*_*x*_ = *R*^*T*^ Σ_*x*_*R* and *G*_*y*_ = *R*^*T*^ Σ_*y*_*R*. The matrix *R* ∈ ℝ^*N* ×*N*^ is orthogonal and Σ_*x*_, Σ_*y*_ are block diagonal, with 2 × 2 skew-symmetric blocks. This decomposition into two-dimensional rotational subspaces matches the structure derived for actionable representations of space [24], and here follows directly from commuting skew-symmetric generators. Assuming *N* is even, the similarity function simplifies to a sum of cosines parameterized by the eigenvalues of the generators. The final cosine similarity evaluates to

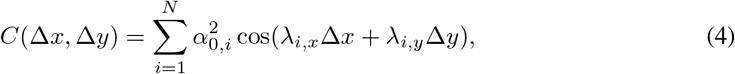

where 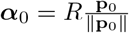 are the imaginary parts of the *i*-th eigenvalues of *G*_*x*_ and *G*_*y*_ (see Appendix F).

### 2.4 Preserving the metric of flat space and grid scale ratios

A desirable property of spatial mapping is the faithful translation of physical Euclidean distances into distances on a neural manifold, which trained models pursue as an objective, either conformal isometry [22, 23] or extrinsic distance preservation [21, 25]. Here, demanding metric preservation (that is, a conformal isometry) of a path-integrating, orthogonal representation yields a closed-form condition on the spectra of the generators (see Appendix E). With the eigenvalues written in polar coordinates as *λ*_*i,x*_ = *k*_*i*_ cos(*ϕ*_*i*_) and *λ*_*i,y*_ = *k*_*i*_ sin(*ϕ*_*i*_), and amplitudes *ρ*_*i*_ = *α*_0,*i*_*k*_*i*_, the representation preserves the flat metric exactly when the weighted phasors at twice the wavevector angles cancel, 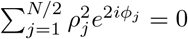. We refer to spectra satisfying this condition as balanced configurations.

The condition is satisfied by partitioning the eigenvalue pairs into modules whose angles form roots of unity on discrete rings in the frequency domain. For a ring of radius *ρ*_*m*_ with symmetry order *M*, the angles are distributed as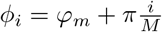, with *i* = 0, 1, …, *M* − 1, and each such module cancels on its own, independently of its orientation *φ*_*m*_. For modules of equal amplitude, roots of unity are also necessary when *M* = 2 or *M* = 3, since two or three equal phasors can only cancel if they are antipodal or form an equilateral triangle. From *M* = 4 onward, irregular balanced configurations exist as well. To visualize the spatial representation produced by a particular symmetry *M*, we note that for orthogonal matrices the entries of the population vector **p** constitute superpositions of 2D plane waves (Appendix C). For a single ring, lower-order symmetries such as *M* = 2 and *M* = 3 produce plane waves oriented 90^°^ and 60^°^ apart, respectively. The combination yields grid-like representations, forming square-type grids for *M* = 2 and hexagonal grids for *M* = 3 (Fig. 2). The radius and orientation of a ring mirror the spacing and orientation of biological grid patterns. One possible criterion for preferring a particular symmetry order *M* is distance preservation over large displacements, which could aid in real-world navigation. Among periodic representations that preserve distances locally, the hexagonal grid (*M* = 3) has the smallest higher-order distortion [25].

**Figure 2:**
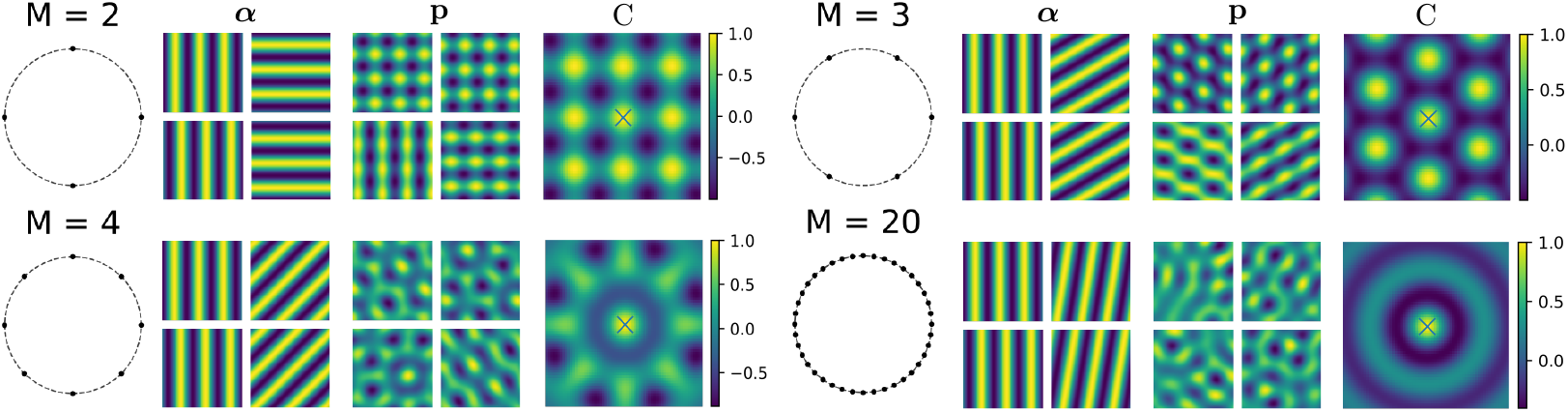
Metric-preserving representations and roots of unity. For each symmetry order *M*, from left to right: the eigenvalue ring (black dots, each root with its conjugate), example plane waves (***α***), example units of the population vector (**p**), and similarity (*C*) relative to the origin (black cross).

Analyzing the similarity function (see Appendix G) reveals that if equally weighted modules with the same spacing have evenly spaced orientations across the range of distinct grid orientations, the representational similarity becomes approximately independent of heading direction over a range of spatial displacements. Furthermore, if the wavenumbers of different modules are proportional to the positive zeros of the Bessel function *J*_0_, the radial component becomes a truncated Fourier-Bessel series. The weight given to each spacing can then be chosen to approximate a range of radially symmetric similarity functions, which may support navigation. Intriguingly, beyond the first, the average of the low-order ratios of successive Bessel function zeros is close to the mean spacing ratio between successive grid modules reported experimentally [34], suggesting a link between the roots-of-unity algebraic structure and the modular organization observed in the entorhinal cortex.

### 2.5 Similarity preservation and context-dependent remapping

Animals encode a variety of non-spatial information, such as room identity or olfactory cues. Unlike spatial position, such information has no intrinsic Euclidean metric to preserve. Instead, we require that similar context signals produce similar representations, while dissimilar contexts result in near-orthogonal ones. Prior studies have shown that such similarity preservation yields localized receptive fields [35], which resemble biological place fields when the inputs are spatial [19].

To preserve similarity rather than the metric, we note that Eq. (4) acts as a discrete approximation of the inverse Fourier transform of a symmetric function with a non-negative Fourier spectrum. Coefficients and eigenvalues can therefore be chosen to approximate a wide class of similarity functions without gradient-based optimization. With a Gaussian target for the spatial similarity, this construction produces localized place-like fields (Appendix H).

In the hippocampal formation, spatial and non-spatial information is encoded conjunctively as part of a unified representation of the environment. When the context changes, the spatial firing patterns can change as well, a phenomenon known as remapping [10, 11]. We extend the exponential map framework to remapping with a context generator *G*_*s*_, which in the simplest case encodes a global scalar context signal *s* in the same way as the spatial coordinates,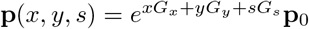. When *G*_*s*_ is skew-symmetric and commutes with *G*_*x*_ and *G*_*y*_, the similarity keeps the form of Eq. (4), with a context term *λ*_*i,s*_Δ*s* added to the argument of each cosine, where *λ*_*i,s*_ is the imaginary part of the *i*-th eigenvalue of *G*_*s*_. To obtain a Gaussian similarity 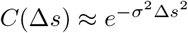 between contexts, we sample the *λ*_*i,s*_ from a normal distribution, the Fourier transform of the target, with uniform coefficients 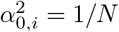 (Appendix I). Dissimilar contexts then become effectively decorrelated.

Across contexts, the representation of a fixed location changes, as in remapping. Within a fixed context, the similarity remains purely spatial, so the geometric structure is the same in every context. Figure 3 illustrates this for a metric-preserving spatial representation combined with a similarity-preserving context generator. The context shift moves and reshapes the fields of individual units, while the lattice periodicity is unchanged.

**Figure 3:**
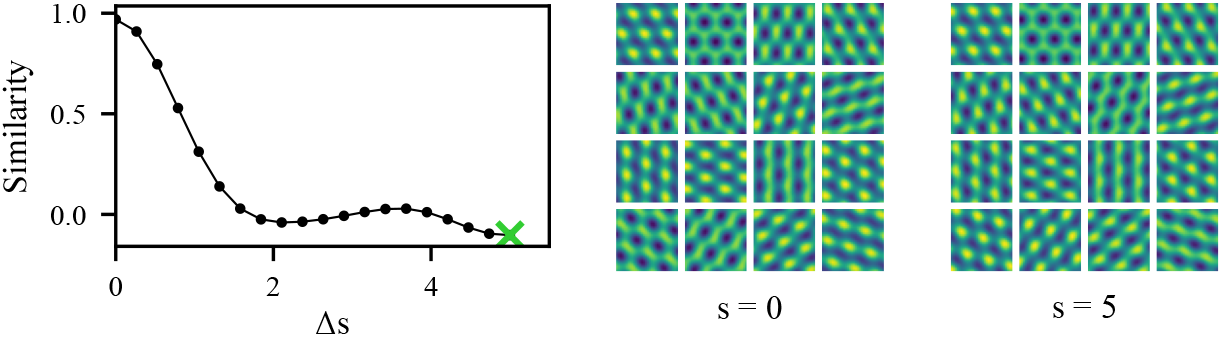
Context-dependent remapping and similarity preservation. Between-context similarity decays rapidly with context separation Δ*s*. Example rate maps show that shifting the context from *s* = 0 to *s* = 5 decorrelates the activity fields, shifting and modulating the fields of individual units while preserving the hexagonal lattice. The population consists of 50 hexagonal sets of six units with distinct orientations, one set per row of rate maps, and *R* mixes within a set, so that the pairwise phase relations within a set are unchanged by the shift (Appendix M). *N* = 300, *M* = 3.

## 3 Biological realization and functional navigation

While derived from abstract algebraic principles, the exponential map framework intersects with established models in theoretical neuroscience. We show that our framework provides a formal connection between the continuous attractor dynamics and oscillatory interference theories. Additionally, we show that it can be fitted to experimental grid and place cell recordings and extrapolates beyond the fitted region (Section 3.2), and that it enables goal-oriented navigation via hyperdimensional binding and bundling.

### 3.1 Relation to attractor dynamics and oscillatory interference

CANN and OI models offer two mechanistic accounts of grid cells, and hybrid models combine them in a single network [36, 37]. The generators of the exponential map relate to both. As matrices, they act like the recurrent weights that move the activity pattern in a CANN, and their spectra describe the oscillators of an OI model.

Consider a recurrent network *τ* **ż**= *U* (**v**)**z** − *MΣ*(**z**) with time constant *τ* . The velocity **v** scales skew-symmetric recurrent weights, *U* (**v**) = *v*_*x*_*U*_*x*_ + *v*_*y*_*U*_*y*_, and the second term, with *M* positive definite and 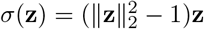, pulls the activity toward unit norm. The state converges to the unit hypersphere, where the second term vanishes and the dynamics reduce to *τ* **ż** = *U* (**v**)**z** (Appendix J.1). If *U*_*x*_ and *U*_*y*_ commute, the state depends only on the current position and not on the path taken to reach it, and is described by the exponential map with *G*_*x*_ = *U*_*x*_*/τ* and *G*_*y*_ = *U*_*y*_*/τ* (Appendix J.2). Since the generators scale as 1*/τ*, the grid spacing is proportional to the network time constant (Appendix J.3).

In CANNs, the symmetric part of the recurrent weights stabilizes the activity pattern, while the skew-symmetric part shifts it [38]. The generators correspond to the skew-symmetric part. The stabilizing term of our network is simpler than in a CANN, as it fixes only the norm of the population vector. Its attractor is therefore the whole hypersphere, of which the spatial code visits a torus, two-dimensional for a single square or hexagonal module. Finally, we demonstrate a local learning rule driven by time-lagged correlations that leaves only the skew-symmetric part of the weights (Appendix J.4), but does not by itself produce commuting generators or the spectra required for a coherent map.

The connection to OI models comes from the spectrum of the generators. In the block-diagonal form of Section 2.3, the population vector decomposes into two-dimensional subspaces, each rotating by a phase *λ*_*i,x*_*x* + *λ*_*i,y*_*y* that grows with the distance traveled along one direction (Appendix C). In OI models, a velocity-controlled oscillator accumulates the same phase relative to a baseline oscillation [13]. Each subspace corresponds to one such oscillator, with the eigenvalues setting its preferred direction and spatial scale (Appendix D). The rate map of a unit is a linear combination of the resulting plane waves, which correspond to the interference bands of OI models, but without a temporal oscillation. Recurrent weights and oscillators are in this sense the same generators, viewed as a matrix and through its spectrum.

### 3.2 Fitting the exponential map to experimental recordings

The framework can also be fitted directly to recorded population data. We fitted generators to the rate maps of 102 grid cells from one module of a rat foraging in an open field [39] and of 103 spatially tuned CA1 place cells of one rat from an open-field dataset [40], with one model unit per recorded cell (Fig. 4). The generators keep the form of Section 2.3, *G*_*x*_ = *R*^*T*^ Σ_*x*_*R* and *G*_*y*_ = *R*^*T*^ Σ_*y*_*R*, with the orthogonal matrix *R* and the wavevectors in Σ_*x*_, Σ_*y*_ as the learnable parameters, so that skew-symmetry and commutativity hold by construction and the loss is the one-step prediction error between neighboring pixels alone (see Appendix K for training details and extended results). Rate maps are generated by transporting the population vector recorded at the arena center.

**Figure 4:**
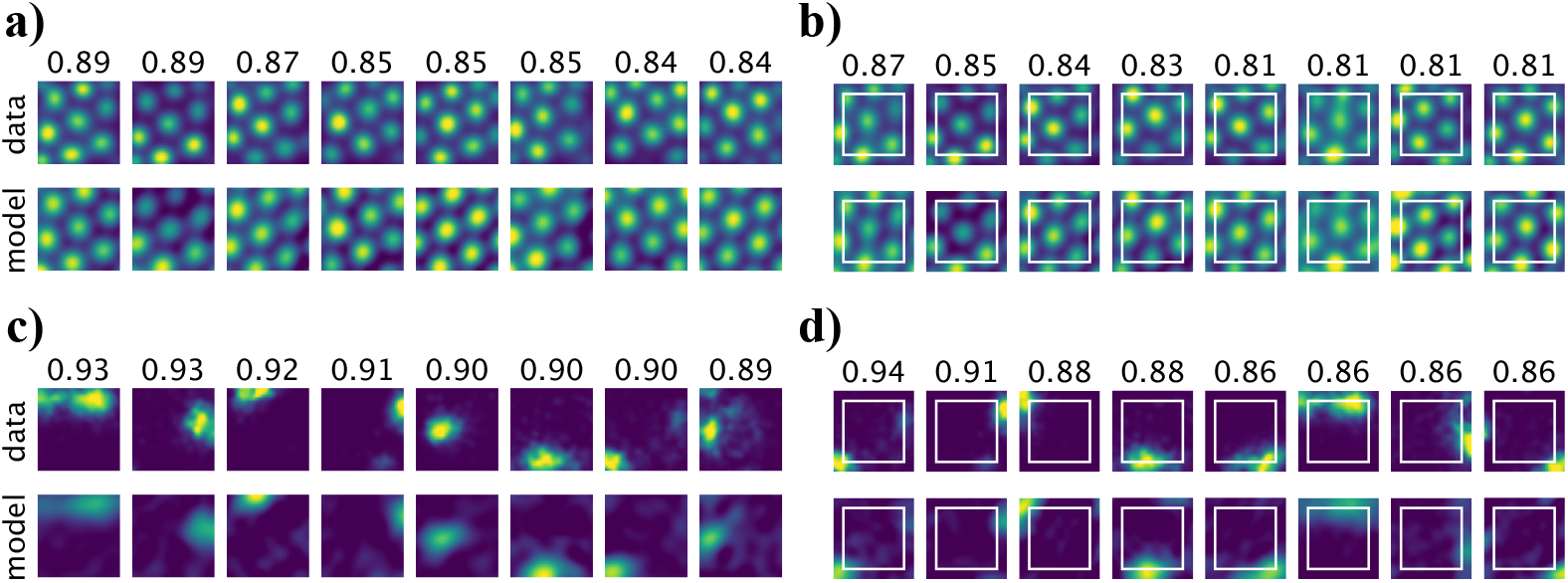
Exponential maps fitted to experimental grid and place cell rate maps. **a)** Rate maps of recorded grid cells [39] (top) and those generated by the fitted model (bottom), for the eight best-matched units, with the correlation given above each pair. **b)** The model is fitted to the central three quarters of the arena side (white square) and the full map is generated from the center. Above each pair is the correlation between data and model over the part of the arena not trained on, for the eight units where it is highest. **c), d)** As a) and b) for CA1 cells [40]. All units are shown in Appendix K.

The generated maps match the data at a median per-unit correlation of 0.76 for grid cells and 0.77 for place cells. To test extrapolation against recorded rate maps, we fitted models to a central square spanning three quarters of the arena side and generated the full map from its center. On the part of the arena the models were not trained on, the generated maps match the data at a median correlation of 0.71 for grid cells and 0.61 for place cells (Appendix K). Fields that extend into the trained square are continued outward. Fields lying entirely outside it are mostly recovered for grid cells and rarely for place cells, as the trained square determines a periodic pattern but not the position of an isolated field. Notably, however, both the grid and the place cell models extend the representation into the part of the arena they were not trained on.

When the arena side is doubled, the generated grid lattices continue periodically beyond the recorded enclosure, and all but one of the generated place cell maps have further place-like fields outside it (Appendix K). No recordings exist there, but CA1 place cells are reported to have multiple, irregularly arranged fields in larger environments [41]. This is also notable, as trained neural network models often struggle to extrapolate beyond the arena they were trained in [17], although recent self-supervised approaches fare better [20].

Grid and place cell recordings are both approximated by the construction derived above, with the data selecting only the wavevectors and the mixing matrix *R*. The fitted representations therefore path integrate exactly and have translation-invariant similarity. The fitted similarity function matches that of the recordings, and for the grid cells it is carried by one hexagonal set of wavevectors (Appendix K).

### 3.3 Goal-oriented navigation via hyperdimensional bundling and binding

The algebraic structure of our spatial representations is deeply tied to hyperdimensional computing [32], natively supporting robust goal-oriented navigation via *bundling* and *binding* (Appendix L). Bundling aggregates representations via element-wise addition into a composite memory vector. Target locations 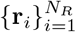 within a specific context *s* are bundled as

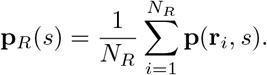

Querying the current location against this memory yields a similarity score

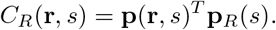

When the pairwise similarities are designed to approximate Gaussians (see Appendix H), this query generates a continuous similarity landscape (a Gaussian mixture) supporting navigation via gradient ascent toward target rewards, **r**_*t*+1_ ← **r**_*t*_ + *η*∇_**r**_*C*_*R*_.

Binding, in contrast, combines vectors into a result that is nearly orthogonal to its constituents. In our formalism, context-dependent remapping functions as an intrinsic binding operation. A context shift applies the orthogonal transformation 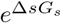, which preserves similarities within a context and makes representations of dissimilar contexts nearly orthogonal (Section 2.5). The memories of several contexts can therefore be bundled into one composite memory, and a query in one context retrieves its own targets while the others largely cancel. An agent can thus switch from foraging to homing in the same arena by changing the context signal *s* and querying the composite memory (Fig. 5).

**Figure 5:**
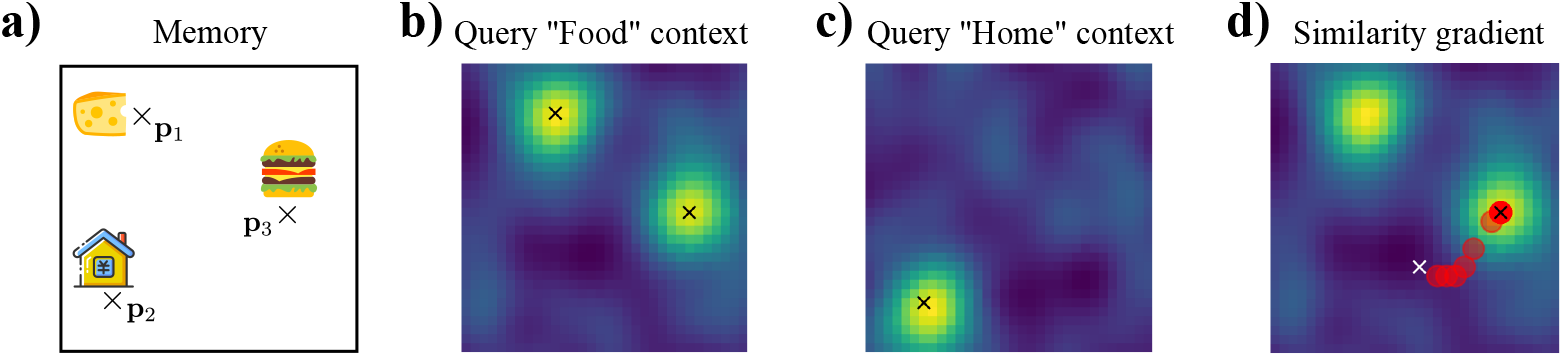
Multi-map, reward-oriented navigation using exponential maps. **a)** A composite memory vector stores the locations of three objects (Cheese, Hamburger, Home) belonging to two distinct contexts (Food and Home). **b)** Querying the memory in the “Food” context retrieves a similarity map that highlights the two food items. **c)** Querying in the “Home” context retrieves the home location. **d)** Gradient ascent on the “Food” similarity map generates a trajectory toward a reward. The similarity function was constructed to be approximately Gaussian, with *N* = 300 units.

### 3.4 Accommodated phenomena and predictions

The framework accommodates several established findings post hoc. First, grid spacing is proportional to the network time constant (Section 3.1), consistent with the dorsoventral gradients of oscillation frequency, h-current time constant and synaptic integration window in medial entorhinal stellate cells [42, 43, 44]. Second, the population activity of a single hexagonal module is confined to a two-dimensional torus (Section 3.1), the topology reported for grid modules [39]. Third, a context shift rotates each two-dimensional rotational subspace by the same phase for all neurons. If subspaces with the same wavevector share a context eigenvalue, the spatial cross-correlogram of any two cells in a module is unchanged by remapping for any mixing matrix *R*, while single-cell fields remain free to deform (Appendix M). This is consistent with the constant phase structure of grid cell ensembles under remapping [11] and with cell-pair relationships conserved despite deformations of single-cell responses [45].

Beyond these, the algebraic structure yields a prediction. Grid modules with evenly distributed orientations at each spacing can jointly approximate radially symmetric similarity functions through a Fourier-Bessel series when the module wavenumbers are proportional to successive zeros of *J*_0_ (Appendix G). The ratio between neighboring module spacings is then not constant but increases toward the coarser modules, following the ratios of successive Bessel zeros. Averaged over the low-order ratios beyond the first, however, it is close to the mean ratio reported experimentally [34].

## 4 Conclusion and limitations

We have introduced an interpretable, exact algebraic formulation of spatial representations, in which generator matrices couple spatial and representational displacement and replace black-box training. Commuting generators guarantee path-independent representations, skew-symmetric generators yield equinorm representations with translationally invariant similarity, and metric preservation reduces to a closed-form balance condition on the generator spectra, which roots of unity satisfy. Despite its mathematical simplicity, the framework supports periodic grid fields and localized place-like fields, and extends to context-dependent remapping through a generator for non-spatial inputs. The framework also relates to attractor and oscillatory interference models of grid cells, supports goal-oriented navigation across contexts, and can be fitted to recorded grid and place cells, whose rate maps it extrapolates beyond the training region.

The approach has several limitations. First, the model assumes open, flat Euclidean environments. Environments with obstacles or curved geometry may call for position-dependent or non-commuting generators. Second, we have only considered the idealized commuting case, and robustness to velocity noise and imperfect generators, as well as the role of sensory corrections at boundaries, remains an open question. Third, our remapping model uses a scalar context signal. Generalizing to high-dimensional structured inputs, such as visual scenes, is an important next step. The algebraic structure naturally supports vector-valued context signals with one generator per component (**s** · **G**_*s*_), so that similar inputs would again produce similar representations. Fourth, the fits to data cover one grid module and one CA1 population, each from a single animal, and extrapolation was tested only within the recorded arena. Fifth, the similarity functions of Section 2.5 are approximated with randomly sampled eigenvalues. The eigenfunctions of the Laplacian are the optimal basis for truncated series of smooth functions [46], which suggests that structured choices of eigenvalues could improve the approximation while favoring periodic solutions. Finally, while geometric considerations fix most parameters, the orthogonal matrix *R* remains a degree of freedom. It strongly influences the tuning curves by mixing the underlying plane waves, but not the similarity function, since for any *R* the initial vector **p**_0_ can be chosen to give the same weights ***α***_0_ in Eq. (4). Outside the fits to data, where *R* is learned, we used randomly sampled orthogonal matrices. Future work should investigate whether biological constraints, such as non-negative firing rates, could inform particular choices of *R*.

## 5 Acknowledgments and disclosures

We thank Frederik Rogge and Anders Malthe-Sørenssen for inspiring discussions in an early stage of the project.

## Author Contributions

Conceptualization: MP, NH, ML. Formal Analysis: MP, NH, ML. Validation: MP, NH, JB, TMS, ML. Writing - Original Draft: MP, NH, ML. Writing - Review & Editing: MP, NH, JB, TMS, ML.

## Code Availability

All code required to reproduce the results and figures in this work is publicly available at https://github.com/bioAI-Oslo/ExponentialRepresentations.

## Use of LLMs

Large language models were used to assist with writing and with the code for the analyses and figures; all derivations, code and results were checked by the authors.

## Appendix

### A Methods

All computational simulations were implemented using the matrix exponential operator in PyTorch [47]. Random orthogonal matrices were generated using the ortho_group module from the SciPy library [48], which samples uniformly from the orthogonal group *O*(*N* ). For experiments involving root-of-unity solutions, the spatial domain was set to 20 × 20 to capture the full periodicity of the representation. For the similarity-preserving experiments, the domains were defined as *s* ∈ [0, 5] for the non-spatial context signal and (*x, y*) ∈ [− 2, 2] for the spatial coordinates, commensurate with the length scale of the target Gaussian similarity function.

### B Commuting generators produce path-independent representations

Since the spatial representation is defined by the exponential map in Eq. (1), we can enforce specific geometric properties by constraining the generators *G*_*x*_ and *G*_*y*_. For example, Schaeffer et al. [20] proposed that representations should be path-independent. The requirement dictates that the representation at a given location must not depend on the trajectory taken to reach it. In the exponential map formalism, the geometric condition is satisfied if the generators commute. To illustrate the property, consider two distinct paths to a point *D* starting from *A*. We define a path *A* →*B*→*D* and an alternate path *A* →*C* →*D*. The resulting representations are generated by composing the transformations for each segment

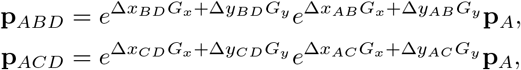

where **p**_*A*_ denotes the representation at *A*. If *G*_*x*_ and *G*_*y*_ commute, their linear combinations also commute. By the Baker-Campbell-Hausdorff (BCH) formula, the composite transformations combine into a single exponential summing the exponents

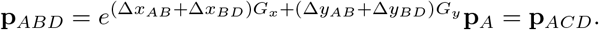

The final state depends only on the net displacement from the initial location, which is identical for both paths.

Conversely, if the generators do not commute, path dependence arises from non-vanishing terms in the BCH expansion. Applying the expansion to the path *A* →*B* →*D*, we define the first displacement as *U* = Δ*x*_*AB*_*G*_*x*_ + Δ*y*_*AB*_*G*_*y*_ and the second displacement as *V* = Δ*x*_*BD*_*G*_*x*_ + Δ*y*_*BD*_*G*_*y*_. Since matrix operators apply to the state vector from right to left, the combined transformation evaluates to

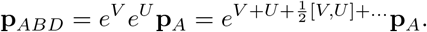

The commutator term [*V, U* ] evaluates to −[*U, V* ] which expands to

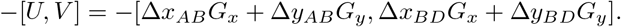

Using the linearity of the commutator and the property [*G*_*x*_, *G*_*y*_] = − [*G*_*y*_, *G*_*x*_], the expansion simplifies to

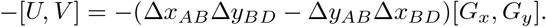

Crucially, if [*G*_*x*_, *G*_*y*_] ≠ 0, the exponent includes a term proportional to the cross product of the path segment displacements. The term geometrically relates to the area enclosed by the path components.

Similarly, for the path *A* →*C* →*D* with alternate operators *W* and *Z*, the commutator [*Z, W* ] introduces different displacement cross-terms. Consequently, the higher-order corrections differ between the two paths, forcing **p**_*ABD*_ ≠ **p**_*ACD*_. Thus, commutativity of the generators is necessary for the representation to depend solely on the net displacement.

### C From generators to plane waves

The structure of the spatial representation defined by Eq. (1) allows for an explicit decomposition of individual cell responses. Assuming a unit-norm origin state **p**_0_ and commuting skew-symmetric generators in block-diagonal form, the dynamics in the rotated basis ***α*** = *R***p** evaluate to

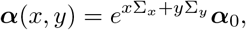

where ***α***_0_ = *R***p**_0_. In the canonical basis, the matrix exponential reduces to a block-diagonal matrix populated by 2 × 2 rotation matrices. Following Dorrell et al. [24], the representation decomposes into independent rotations within distinct two-dimensional subspaces. For the *i*-th block, the update rule evaluates to

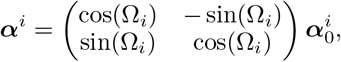

where ***α***^*i*^ ∈ ℝ^2^ represents the state vector within the *i*-th subspace. The rotation angle Ω_*i*_ = *xλ*_*i,x*_ + *yλ*_*i,y*_ couples the spatial displacement to the generator eigenvalues.

Letting 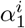 and 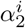 denote the two components of the *i*-th block (corresponding to the conjugate eigenvalue pair), the matrix multiplication yields

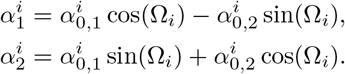

By applying the harmonic addition theorem, the components can be rewritten as phase-shifted sinusoids

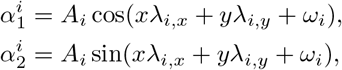

where the amplitude is 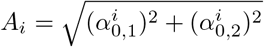 and the phase is *ω*_*i*_ = arctan 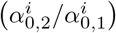.

The equations demonstrate that each canonical component acts exactly as a two-dimensional plane wave with orientation and frequency determined by *λ*_*i,x*_ and *λ*_*i,y*_, alongside a phase shift *ω*_*i*_ along the wave direction. Since the observed neural representation is given by **p** = *R*^*T*^ ***α***, the abstract firing rate of each neuron consists of a linear superposition of the plane waves.

### D Mathematical connection to oscillatory interference

The algebraic framework established by the exponential map provides a direct mathematical formalization of the spatial components of oscillatory interference (OI) models.

In the classical OI model [13], grid cell firing arises from the interference between a somatic baseline oscillation of angular frequency *ω*_*s*_ and multiple dendritic velocity-controlled oscillators (VCOs). The angular frequency of a dendritic VCO, *ω*_*d,i*_, is modulated by the animal’s speed *v* and its heading *ϕ* relative to the VCO’s preferred direction *ϕ*_*d,i*_ according to the phase equation

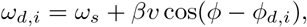

where *β* is a spatial scaling constant. Integrating the frequency over time yields the phase of the dendritic oscillator

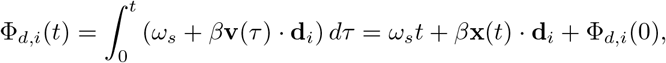

where **v**(*t*) is the velocity vector, **x**(*t*) is the position vector, and **d**_*i*_ is the unit vector for the preferred direction.

The somatic and dendritic membrane potentials summate, generating a predictable interference pattern. For equal amplitudes *A*, the interference between the somatic and *i*-th dendritic oscillator evaluates to

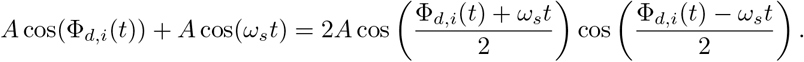

The neuron fires at the peaks of the high-frequency carrier wave, modulated by the lower-frequency spatial envelope. Substituting the integrated phase, the envelope evaluates to

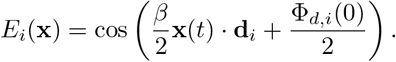

The interference pattern forms a one-dimensional spatial plane wave. The product of multiple interference patterns generates the two-dimensional hexagonal grid.

The mechanism mathematically mirrors the spectral decomposition of the exponential map model derived in Appendix C. For skew-symmetric commuting generators *G*_*x*_ and *G*_*y*_, the representation decomposes into independent two-dimensional rotational subspaces. The components of the *i*-th subspace evolve according to

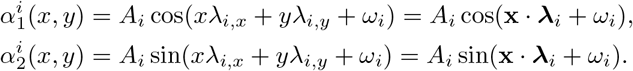

The components have the form of the spatial envelopes of the OI model, with the correspondence

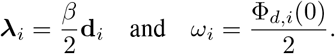

The generator eigenvalues *λ*_*i,x*_ and *λ*_*i,y*_ thus play the role of the wavevector components set by the VCO preferred directions 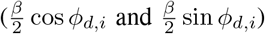. The correspondence is between the signed envelopes. The firing of an OI neuron follows the magnitude of its envelope, so its bands recur at wavenumber *β*, and the OI model combines the bands of several VCOs multiplicatively, whereas the exponential map superposes plane waves linearly through *R*.

### E Metric preservation

Consider the representation along a parametrized trajectory **r**(*t*) = (*x*(*t*), *y*(*t*)) defined by

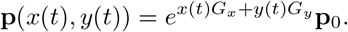

The length of the trajectory in the representational space is calculated by the path integral of the line element *ds* = *∥d***p***∥*_2_. By the chain rule, the differential change in the representation is

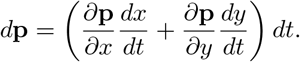

The trajectory length *L* can be expressed in terms of the induced metric *g*_*ij*_ as

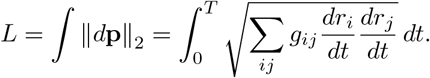

Comparing the length with the squared line element, we evaluate the induced metric *g* as

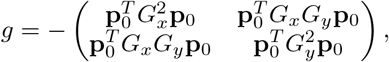

where we use the identity 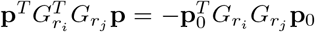 derived from the skew-symmetry of the generator matrices.

We simplify the metric using the block-diagonal decomposition *G* = *R*^*T*^ Σ*R*. Since Σ contains 2 × 2 skew-symmetric blocks, 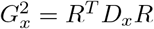, where *D*_*x*_ is a diagonal matrix containing the squared imaginary part of a given eigenvalue 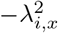. The mixed term *G*_*x*_*G*_*y*_ similarly diagonalizes to entries −*λ*_*i,x*_*λ*_*i,y*_. Letting ***α***_0_ = *R***p**_0_, the metric simplifies to

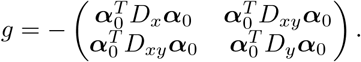

Substituting the metric back into the path length integral yields

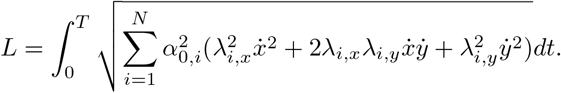

To preserve the flat Euclidean metric, we require the off-diagonal terms to vanish and the diagonal terms to be equal

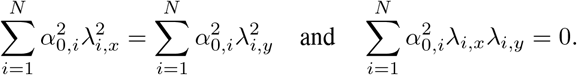

Introducing polar coordinates for the eigenvalues *λ*_*i,x*_ = *k*_*i*_ cos *ϕ*_*i*_ and *λ*_*i,y*_ = *k*_*i*_ sin *ϕ*_*i*_ and defining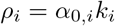, the conditions evaluate to

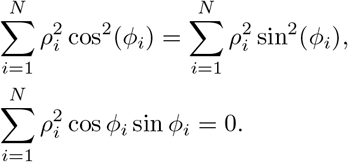

Using trigonometric identities, the system simplifies to requiring that the weighted sum of phasors vanishes at double the angle

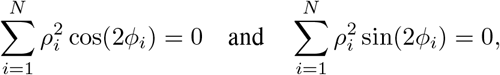

which is mathematically equivalent to the complex condition

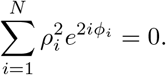

Since the two components of a pair share the wavevector 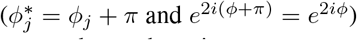, the sum reduces to one over the *N/*2 pairs, with the weight 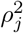 summed over the pair.

Assuming equal weighting *ρ*_*i*_ = *ρ*, the condition reduces to finding a set of angles such that

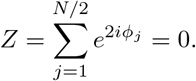

The condition is satisfied if the eigenvalue pairs are partitioned into subsets or modules where the spatial phases *ϕ*_*i*_ form a collection of roots of unity that sums to zero. Specifically, for a single module of symmetry order *M*, we require *M* eigenvalue pairs distributed uniformly on the circle

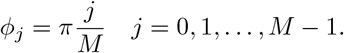

Consequently, the full high-dimensional representation can be constructed as a linear combination of the sets. For a system with multiple modules *k* = 1 … *K*, each with radius *ρ*_*k*_, symmetry *M*_*k*_ and orientation *φ*_*k*_, the total sum vanishes if each module vanishes individually

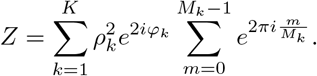

In other words, for each radius *ρ*, there can be multiple rotated sets of roots of unity, each possessing an independent rotational symmetry.

Roots of unity are a sufficient construction rather than the only solution. For a single module of equal amplitude, two phasors cancel only if they are antipodal (*M* = 2, wavevectors 90^°^ apart) and three only if they form an equilateral triangle (*M* = 3, wavevectors 60^°^ apart), so the roots-of-unity structure is necessary in these minimal cases. From *M* = 4 onward, irregular solutions exist, for example the wavevector angles {0^°^, 33^°^, 90^°^, 123^°^}, whose doubled angles {0^°^, 66^°^, 180^°^, 246^°^} cancel pairwise. Unequal amplitudes admit further solutions. Selecting among balanced configurations therefore requires additional criteria, such as distance preservation over larger displacements [25].

A comparison with the explicit form of the representation in Appendix C reveals a striking parallel to the modular organization of grid cells in the medial entorhinal cortex [2, 34], which are similarly organized into modules of a common spacing (*ρ*) and orientation (*φ*).

### F Similarity function derivation

We derive the explicit form of the representational similarity starting from Eq. (3). Because real skew-symmetric matrices commute with their transposes (*GG*^*T*^ = *G*^*T*^ *G*), the matrices are strictly normal and form orthogonal exponentials. Assuming the generators also commute with each other, we apply the simultaneous block-diagonal decomposition

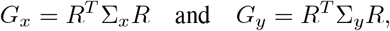

where *R* is a shared orthogonal matrix. Using the identity 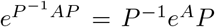, we rewrite the similarity expression as

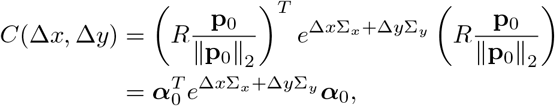

where we define the rotated unit vector 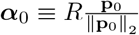 . Note that 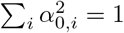 due to the orthogonality of *R*

The exponent matrix Ω = Δ*x*Σ_*x*_ + Δ*y*Σ_*y*_ retains the block-diagonal and skew-symmetric structure of the generators. In the power series expansion of the matrix exponential, even powers result in diagonal matrices while odd powers remain skew-symmetric. Since a quadratic form **x**^*T*^ *A***x** vanishes identically for any skew-symmetric matrix *A*, odd terms do not contribute. The expansion therefore reduces strictly to a sum over even powers

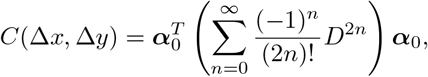

where *D* is a diagonal matrix with entries *θ*_*i*_ = *λ*_*i,x*_Δ*x* + *λ*_*i,y*_Δ*y* and *λ*_*i*,_ denotes the imaginary part of the corresponding eigenvalue. Recognizing the Taylor series for the cosine function, the matrix sum converges to a diagonal matrix with entries cos(*θ*_*i*_). Consequently, the similarity simplifies analytically to

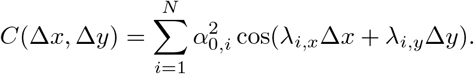

### G Designing spatial similarity functions

We established that the similarity function Eq. (4) takes the general form of a weighted sum of cosines. To understand the structure of the resulting similarity function, we rewrite the expression in polar coordinates where *x* = *r* cos *θ* and *y* = *r* sin *θ* yielding

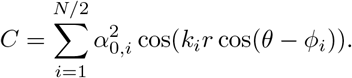

Here, *i* indexes wavevector pairs, and 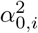 is shorthand for the sum of the two corresponding squared components of the normalized initial vector, so 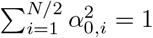 . Applying the Jacobi-Anger expansion evaluates to

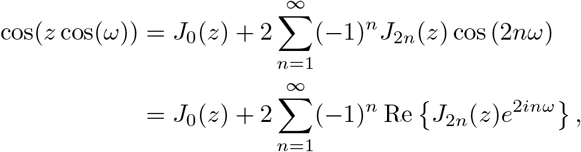

where *J*_*n*_(*z*) is the *n*-th Bessel function of the first kind. Substituting the expansion into the similarity function yields

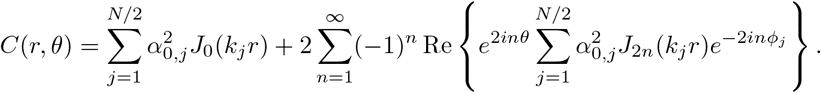

The expansion decomposes the similarity into a purely radial component (the *J*_0_ term) and a mixed term dependent on heading direction *θ*.

Further simplification requires additional structure beyond the general metric-preservation condition. We therefore choose equally weighted roots-of-unity constellations on discrete rings in wavevector space. These are sufficient for metric preservation and, for an individually metric-preserving module with equal weights and a common wavenumber, are the only configurations when *M* = 2 or *M* = 3. These minimal configurations correspond to the wavevector geometries of square and hexagonal grids, respectively. For *K* rings with wavenumbers *k*_*a*_, symmetry orders *M*_*a*_ and orientations *φ*_*a*_, the inner phasor sum becomes

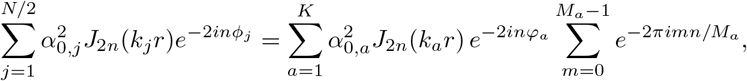

where the geometric series 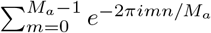 vanishes unless *n* is a multiple of the symmetry order *M*_*a*_. Consequently, angular dependence only arises at harmonic orders *n* = *ℓM*_*a*_. For large *M*, the similarity function becomes approximately isotropic (independent of heading direction).

Angular dependence can then be suppressed even for the low symmetry orders *M* = 2 and *M* = 3 by combining modules with different orientations. At each wavenumber *k*_*a*_, we include *L* equally weighted copies of the same *M* -axis module, with orientation offsets *φ*_*l*_ = *πl/*(*ML*). These orientations evenly subdivide the module’s fundamental orientation interval *π/M*, while each constituent module retains its square or hexagonal grid geometry. The inner phasor sum then becomes

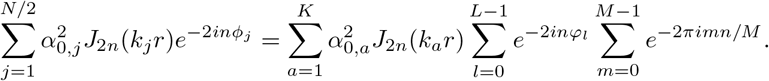

Since each weight 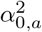 occurs *ML* times, the normalization becomes 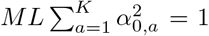. The within-module sum restricts *n* to multiples of *M* . Writing *n* = *qM*, the orientation sum then vanishes unless *q* is a multiple of *L*. Angular terms therefore survive only at *n* = *ℓML*. The full similarity function evaluates to

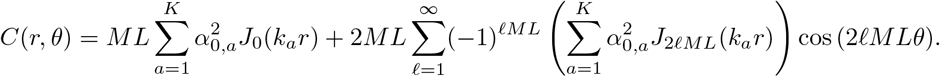

The magnitude of the angular terms is governed by the high-order Bessel functions. For small arguments *z*, the Bessel function of order *γ* behaves according to

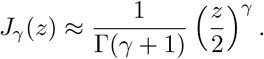

Here, the order is *γ* = 2*ℓML*. For large *ML*, the high-order terms decay rapidly near the origin. Consequently, for a relatively large range of displacements *r*, the angular terms become negligible and the similarity evaluates effectively to its radial component

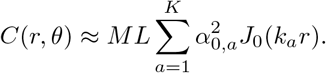

Grid modules may therefore provide a means of suppressing angular dependence: appropriately distributed module orientations allow square or hexagonal grid structure within modules to support approximately isotropic similarity at the population level. Also notable is the fact that choosing *k*_*a*_ = *j*_0,*a*_*/r*_max_, where *j*_0,*a*_ is the *a*-th positive zero of *J*_0_, makes the radial component a truncated Fourier-Bessel series on 0 ≤ *r* ≤ *r*_max_. Thus, modules of grid cells with relative orientations can approximate a range of radial similarity functions by choosing appropriate non-negative coefficients, linking the spatial scales of the modules to the domain of approximation.

In a Fourier-Bessel series the wavenumbers *k*_*a*_ are proportional to the successive zeros of *J*_0_, so the spacing of module *a* is inversely proportional to the *a*-th zero and the ratio between neighboring spacings equals the ratio of successive zeros. These ratios are 2.30, 1.57, 1.36 and 1.27 for the first five zeros and decrease slowly toward one with increasing index (Fig. 6), so the spacing ratio grows with grid scale rather than staying constant. Approximately constant mean grid scale ratios ranging from about 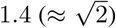 to 1.7 have been reported [34, 49], alongside theoretical accounts linking module scaling to spatial decoding [50]. Our construction suggests that these ratios may also arise from the approximation of a radial similarity kernel in the chosen Fourier-Bessel basis. Finally, we note that while the ratio determines relative scaling, the Fourier-Bessel series inherently defines an absolute length scale related to the domain of approximation. The mathematical relationship suggests that the absolute grid scale may be set by the size of the region the animal needs to encode reliably.

**Figure 6:**
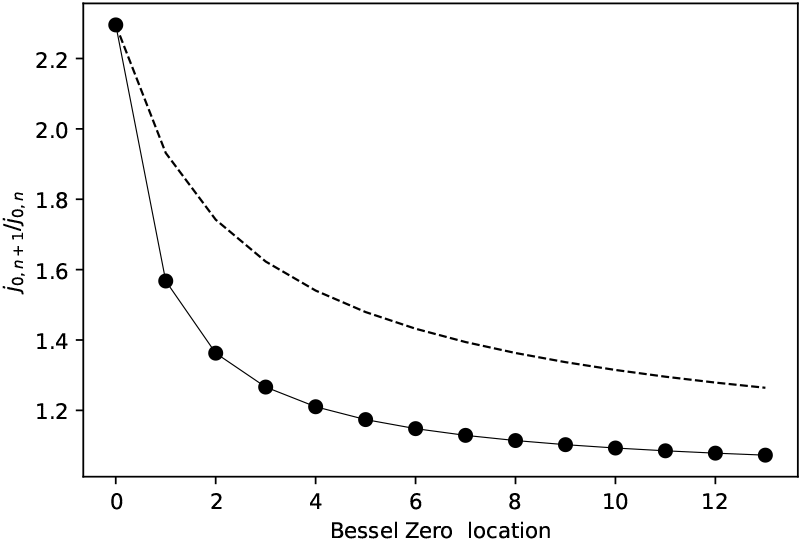
Bessel function zeros and grid scale ratios. Ratio of subsequent zeros of the Bessel function *J*_0_ (solid line with circular markers), alongside a cumulative average (dashed line). Note that the x-axis represents the Bessel zero index.

Suppression of angular dependence also has a consequence for modules with similar grid spacing but different orientations. Their orientations should be approximately evenly distributed over the grid’s rotational symmetry interval. For hexagonal grids, this gives orientation differences of approximately 60^°^*/L* between successive orientations among *L* modules with the same spacing. For example, four such modules should have orientations staggered by approximately 15^°^.

### H Similarity-preserving spatial representations

By relaxing the strict requirement for metric preservation, we can explore the more general case of representations designed to preserve similarity structure.

Following the approach outlined in Appendix I, we can approximate a range of non-negative similarity functions (with non-negative Fourier spectra), by sampling generator eigenvalues according to the target function’s Fourier transform. Notably, this allows us to generate spatial representations with certain similarity functions without any gradient based optimization. As in the contextual case described in Appendix I we can consider the case where generator eigenvalues are sampled from a normal distribution, but in the purely spatial case. This parameterization yields an approximate Gaussian similarity function.

To show that this approach indeed yields meaningful, purely spatial representations, we simulated a population of *N* = 256 units with generator eigenvalues for *G*_*x*_ and *G*_*y*_ sampled from *N*(0, 2). The results are shown in Fig. 7. Unlike the periodic grid-like patterns characteristic of low-order roots-of-unity solutions representing metric preservation, the units exhibit heterogeneous and spatially localized place-like fields, reminiscent of hippocampal place cells [1]. Given that place cells are known to encode both spatial and non-spatial cues such as olfactory context [51], the observation suggests that the spatial model defined in Eq. (1) could be naturally extended to model conjunctive representations of space and context, consistent with recent theoretical proposals [19].

**Figure 7:**
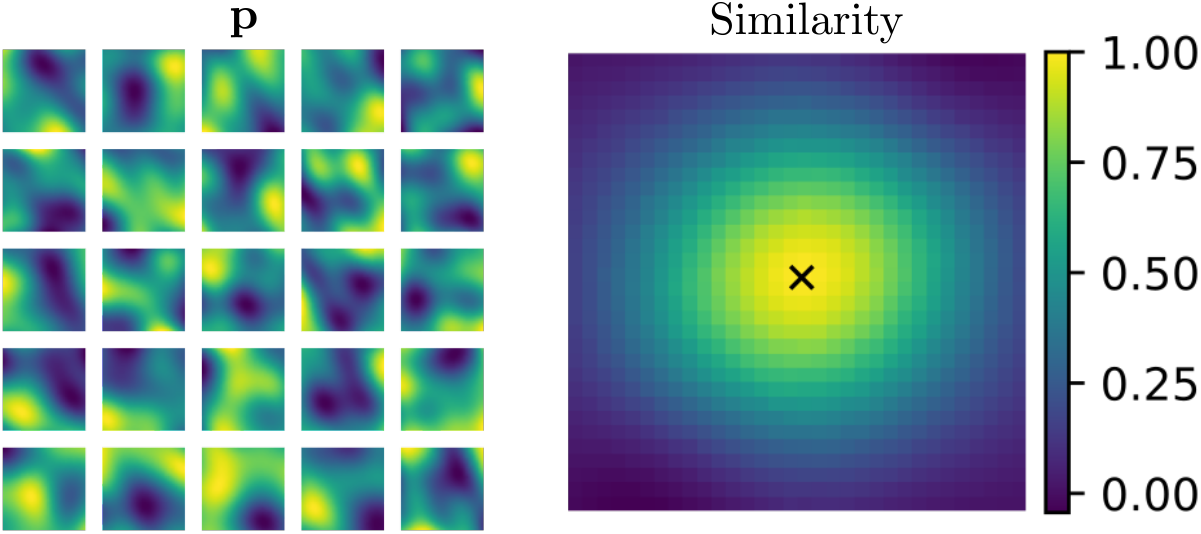
Emergence of localized place-like fields from similarity preservation. Left. Example rate maps for a model where generator eigenvalues are sampled from a normal distribution, resulting in an approximately Gaussian similarity function. **Right**. The resulting spatial similarity function relative to the origin.

The Gaussian similarity function serves as a minimal proof-of-concept demonstrating similarity preservation. The formulation does not yet capture the full statistical complexity, overdispersion or sparsity of empirical place cell firing.Future work must refine the target similarity functions to capture these empirical complexities.

### I Similarity-preserving context representations

By extending similarity-preservation to include non-spatial signals, we can model spatial representations that conjunctively encode context signals. In particular, we can enact contextual similarity preservation by orthogonal transformations that ensure that shifting the context *s* decorrelates the spatial map. This decorrelation allows a single neural population to multiplex distinct reward environments without catastrophic interference, mimicking biological remapping [11]. In the terminology of hyperdimensional computing [32], the contextual decorrelation functions mathematically as an intrinsic *binding* operation.

To implement the mechanism algebraically, we introduce a generator coupled to a scalar context signal *G*_*s*_, and fixed spatial coordinates. In this case, we observe that the similarity function evaluates to

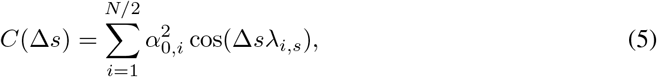

where *λ*_*i,s*_ denotes the imaginary part of the *i*-th eigenvalue of *G*_*s*_ (assuming fixed spatial location). Since the expression is derived from the cosine similarity, the function is strictly bounded to the interval [ −1, 1]. To achieve similarity preservation, we seek a profile *C*(Δ*s*) that decays with increasing Δ*s* towards a baseline level where inputs are deemed dissimilar. We can approximate a wide class of functions by recognizing that Eq. (5) is effectively a cosine series with non-negative coefficients. Specifically, the series operates as a discrete approximation of the inverse Fourier transform of a symmetric function with a non-negative Fourier spectrum

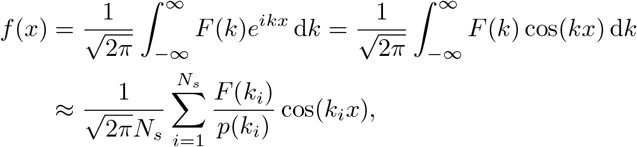

where the summation represents a Monte Carlo estimate of the integral using importance sampling with *N*_*s*_ = *N/*2 samples drawn from probability density *p*(*k*_*i*_).

While several functions meet the specified criteria, an important example is the Gaussian function. The continuous Fourier transform of a target function 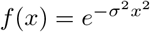 evaluates to

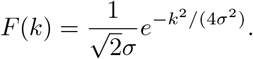

If we sample the eigenvalues *λ*_*i,s*_ from a Gaussian distribution N(0, 2*σ*^2^) with density

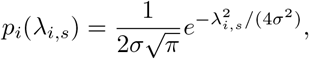

the ratio *F* (*λ*_*i,s*_)*/p*_*i*_(*λ*_*i,s*_) evaluates exactly to 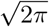. The series coefficients then simplify perfectly to

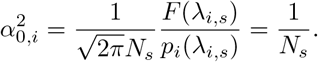

The constant coefficient requirement ensures that the sum approximates the target similarity

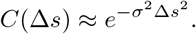

The approximation ensures that similar contexts are highly correlated while dissimilar contexts become decorrelated. Going forward, we set *σ* = 1. Notice that the exact result requires us to select block amplitudes 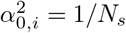, which is achieved if the origin state is initialized as 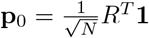 to distribute power equally across all *N* dimensions.

To demonstrate the remapping behavior, we instantiated a metric-preserving spatial representation of 50 hexagonal sets on one ring, each rotated by *π/*150 relative to the previous one, and extended it to encode a non-spatial signal *s* with one context eigenvalue per subspace. The mixing matrix *R* is block diagonal with one 6 × 6 orthogonal block per set, so that each unit draws on the three wavevectors of one set. Many sets are needed for the Monte Carlo estimate to approximate the Gaussian. Since no two sets share a wavevector, every wavevector carries a single context eigenvalue, and the pairwise phase relations within a set are unchanged by the context shift (Appendix M). Sets whose orientations differ by a few degrees cannot be told apart within the arena, so this invariance holds within a set and not between neighboring sets. The resulting between-context similarity is shown in Fig. 3. Similarities decay with increasing context dissimilarity, and the rate maps of unit activity shift between contexts, reproducing remapping dynamics [11]. Crucially, spatial similarities are preserved for any fixed *s*, maintaining the grid-like structure derived in the previous section.

### J Biological interpretation of attractor networks

The exponential map framework provides a rigorous algebraic description of spatial representations. Here we demonstrate that the framework is not merely an abstraction but emerges as the on-manifold dynamics of a gain-modulated continuous attractor neural network (CANN).

#### J.1 Continuous attractor formulation

We consider the general dynamics of a recurrent neural network [38, 52] governed by the equation

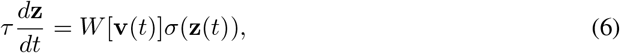

where **z** *∈* ℝ ^*N*^ is the neural state vector and *τ* is the effective time constant. Here *W* [**v**] represents the effective state-transition matrix modulated by an external input **v** such as a 2D velocity vector and *σ* is a generalized non-linear function. The formulation subsumes standard leaky-integrator models (for example *τ* **ż**= −**z** + *Jϕ*(**z**)) if

We investigate the case where network activity flows toward a low-dimensional manifold *ℳ*, specifically a unit hypersphere, consistent with the orthogonal transformations derived in our algebraic framework. To ensure stable convergence without singularities we formalize the effective network dynamics as

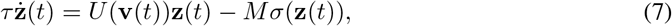

where *U* (**v**) is a velocity-dependent skew-symmetric matrix, *M* is a positive definite matrix and the activation function is defined as 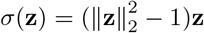.

To verify that the system forms a hypersphere attractor we analyze the Lyapunov energy function

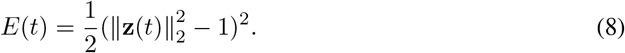

The energy function quantifies the deviation of the state from the manifold surface. The time derivative of the energy evaluates to 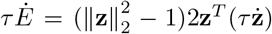. Substituting the temporal derivative from Eq. (7) yields

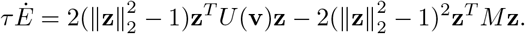

Because *U* (**v**) is defined as a skew-symmetric matrix, the quadratic form **z**^*T*^ *U* (**v**)**z** vanishes identically. The energy derivative simplifies perfectly to

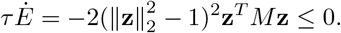

Because *M* is positive definite, the energy strictly decreases for any state off the manifold (*Ė <* 0 for **z** ≠ **0**). The non-increasing property guarantees stable flow toward the hypersphere without requiring a predefined toroidal topology. The attractor is the whole hypersphere, not a position on it. The dynamics correct the norm of the state and leave its position on the sphere free, so an error along the manifold is not corrected.

The decomposition reveals two distinct functional components. The non-linear term (scaled by *M* ) enforces the attractor dynamics normal to the manifold while the linear term (scaled by *U* ) drives transport along the manifold. Whenever the state is on the manifold **z**^*\**^ *∈ ℳ*, the normal-acting attractor contribution vanishes (*σ*(**z**^*\**^) = 0). The macroscopic on-manifold dynamics then reduce to pure transport

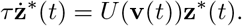

For constant velocity inputs the linear system possesses the exact solution

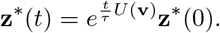

Thus, the exponential map exactly describes the trajectory of the neural state on the continuous attractor manifold.

#### J.2 Connection to the path-integrating exponential map

By defining the asymmetric shift operator *U* (**v**(*t*)) = *v*_*x*_(*t*)*U*_*x*_ + *v*_*y*_(*t*)*U*_*y*_, the solution to the time-varying matrix differential equation involves the time-ordered exponential (*T* ) given by

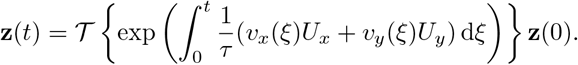

Mathematically the time-ordered exponential expands into the infinite series known as the Magnus expansion [53]. The series includes integral terms involving nested commutators of the operators at different time points such as [*U* (**v**(*ξ*_1_)), *U* (**v**(*ξ*_2_))].

For the system to perform exact path integration, the resulting population vector must depend solely on the net accumulated displacement independent of the specific velocity history or trajectory taken. Path independence requires the time-ordered exponential to reduce to a standard matrix exponential of the integrated inputs. The simplification occurs strictly if the commutator terms in the Magnus expansion vanish, imposing the algebraic constraint that the effective skew-symmetric matrices must commute so that [*U*_*x*_, *U*_*y*_] = 0.

Under the commutative condition we can integrate the velocity inputs directly. Identifying the spatial coordinates as 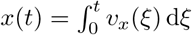 and 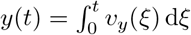 we recover the exponential map model

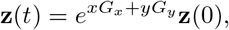

where the generator matrices are identified physically as 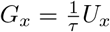 and 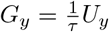.

#### J.3 Membrane dynamics and grid scale hierarchies

The identification of the generator matrices as 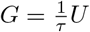 implies an intrinsic physical coupling between the network time constant *τ* and the spatial scale of the resulting representation. As established in Appendix C, the activity of any unit decomposes into a superposition of plane waves. The phase angle φ_*i*_ for the *i*-th subspace is given by the equation

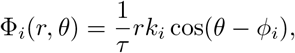

where *r* is spatial displacement, *θ* is heading direction, *k*_*i*_ is the intrinsic frequency and *ϕ*_*i*_ is the orientation of the wave. Considering motion along the wave propagation direction (*θ* = *ϕ*_*i*_), the fundamental spatial period Λ_*i*_ is determined by the condition φ_*i*_(Λ_*i*_, *ϕ*_*i*_) = 2*π*. Solving for Λ_*i*_ yields

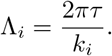

For a hexagonal grid cell representation (*M* = 3), the global grid pattern is constructed from the interference of three such plane waves. Consequently the grid spacing *λ*_grid_ must also scale linearly with the time constant according to

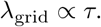

The algebraic derivation mirrors the topographic organization of the medial entorhinal cortex (MEC), where the frequency of subthreshold oscillations [42], the time constant of the h current [43] and the window for synaptic integration [44] in layer II stellate cells vary systematically along the dorsal-ventral axis, in step with the expansion of the grid scale. The framework thus accommodates this gradient and offers a candidate mechanism for it (Section 3.4).

#### J.4 Emergence of skew-symmetric connectivity via local learning

We investigate a biologically plausible local learning rule capable of driving the weight matrix *W* toward skew-symmetry. Motivated by the requirement that the symmetric component of the connectivity must vanish to preserve the manifold energy, we propose the update rule

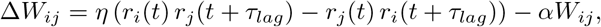

where *r*_*i*_ and *r*_*j*_ denote pre- and postsynaptic activities, *τ*_*lag*_ represents a small temporal lag, *η* is the learning rate and *α* is a local decay term.

Decomposing the weight matrix into symmetric (*S* = *W* + *W*^*T*^ ) and antisymmetric (*A* = *W* − *W*^*T*^ ) components reveals distinct evolutionary dynamics. Since the Hebbian term *H*_*ij*_ = *r*_*i*_(*t*)*r*_*j*_(*t* + *τ*_*lag*_) − *r*_*j*_(*t*)*r*_*i*_(*t* + *τ*_*lag*_) is inherently antisymmetric (*H*_*ij*_ = − *H*_*ji*_), the term contributes zero to the update of the symmetric component. Consequently the dynamics of *S* are governed solely by the decay term

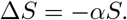

The isolated decay term ensures that any initial symmetric connectivity decays asymptotically to zero. Conversely the antisymmetric component is reinforced by the Hebbian term

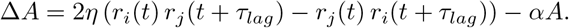

Thus *A* is driven by the time-lagged anti-correlated activity patterns while being stabilized by the decay *α*. In the steady state the antisymmetric part converges to

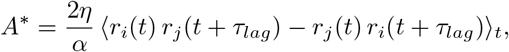

where the brackets denote a time average. The steady state implies that the learned skew-symmetric connectivity is determined by the statistics of the time-lagged antisymmetric correlations in the network activity.

To validate the mechanism we simulated the training of a 32 × 32 weight matrix. We initialized the weights from a normal distribution and drew the rates at every step independently from a normal distribution, with *α* = 0.001 and *η* = 0.01. The evolution of the weights was tracked over 5000 iterations. The results shown in Fig. 8 demonstrate that the magnitude of the symmetric component decays to zero while the skew-symmetric component persists. The persistence shows that the proposed rule filters out symmetric connectivity. For independent rates the time average in *A*^*\**^ falls as 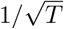, so the persisting component reflects the fluctuations of the correlations rather than a fixed steady state. When the rates are instead transported by a skew-symmetric generator Ω over a small step 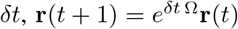 the average converges and the learned *A* matches *A*^*\**^ at a correlation of 0.96, against 0.66 for the independent rates. While the learning rule effectively yields a general skew-symmetric matrix based on input correlations, the rule does not strictly guarantee convergence to the specific commuting generators required for coherent spatial mapping. Future work must investigate how additional constraints or topological priors guide the synaptic dynamics toward exact commutativity.

**Figure 8:**
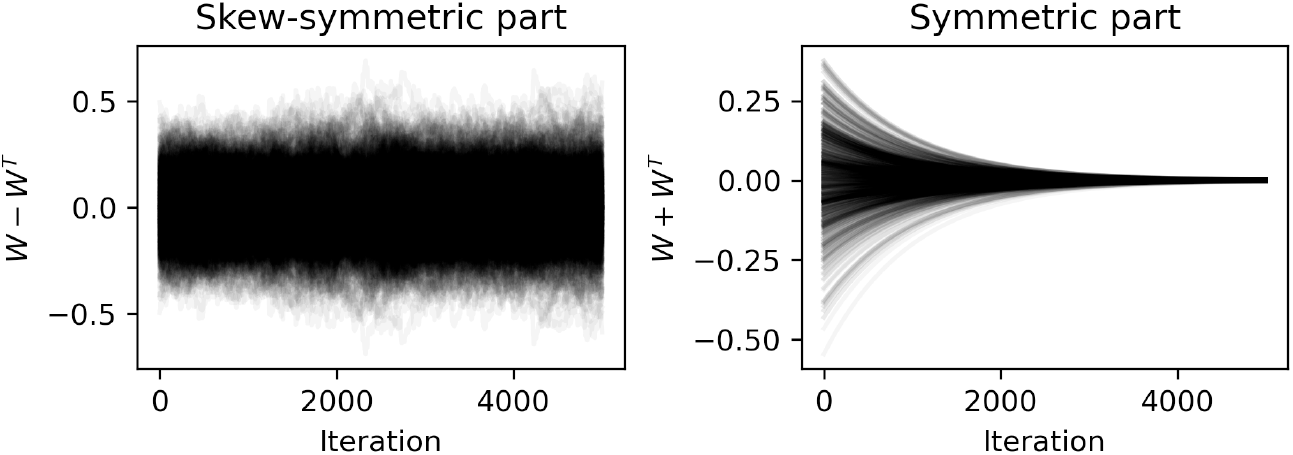
Emergence of skew-symmetric connectivity via local learning. Evolution of weight matrix components under the anti-symmetric Hebbian learning rule. The unscaled symmetric component (*S* = *W* + *W*^*T*^, right panel) vanishes over time, while the unscaled skew-symmetric component (*A* = *W* − *W*^*T*^, left panel) persists.

### K Generative modeling of experimental data

This appendix gives the data, the fitting procedure and the evaluation behind Section 3.2, and shows every fitted unit.

#### Data

Grid cells are 102 cells of one grid module of one rat (module R1, open-field session of day 1) published with [39], pure and conjunctive grid cells alike, each a 32 × 32 rate map over the arena. Place cells come from the open-field CA1 dataset of [40], rats foraging in a 150 × 150 cm arena, published alongside [54], whose preprocessing and unit criteria we follow. Head position was smoothed with a Gaussian of three samples, frames with running speed outside 6 to 60 cm/s were discarded, and spike counts and occupancy were binned at 32 × 32 over the occupied extent of each session (about 4.3 cm per bin), smoothed with a Gaussian of one bin and divided. Clusters nested within a larger cluster of the same session, sharing more than half their spikes, were treated as duplicates, and units were kept if their mean rate over the session lay between 1 and 10 Hz as in the source paper. Since the fit models spatial structure, we further required a spatial information of at least 0.5 bits per spike, and we took every such unit of one rat, so that one model describes one animal’s place code. Few units were recorded in any one session, so the population is pooled over the rat’s 37 sessions and its population vectors were never observed simultaneously. Tetrodes stay in place across days, so units on the same tetrode recorded on different days whose rate maps correlate above 0.9 were treated as one cell, of which the one with the highest spatial information was kept. This removed 4 of 107 units and left 103, of which 47 have a single place field and 56 several, a field being a contiguous region above 20% of the peak rate covering at least nine bins.

#### Model and fitting

The population vector **p**(**r**) ∈ ℝ*N* is the vector of recorded rates at position **r**, with one model unit per recorded cell, *N* = 102 for grid cells and *N* = 103 for place cells. The generators are parameterized as in Section 2.3, *G*_*x*_ = *R*^*T*^ Σ_*x*_*R* and *G*_*y*_ = *R*^*T*^ Σ_*y*_*R*, where Σ_*x*_ and Σ_*y*_ are block diagonal with 2 × 2 skew-symmetric blocks holding one wavevector (*λ*_*i,x*_, *λ*_*i,y*_) per block, and 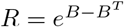 is orthogonal for any matrix *B*. The learnable parameters are *B* and the ⌊*N/*2⌋ wavevectors, so the generators are skew-symmetric and commute for every parameter value. For odd *N* the zero eigenvalue of each generator leaves one direction of population space invariant. Transport is evaluated in closed form as the block rotations of Appendix C, each pair of coordinates in the eigenbasis rotating by Ω_*i*_ = *δx λ*_*i,x*_ + *δy λ*_*i,y*_, which avoids the matrix exponential during training. The loss is the one-step prediction error,

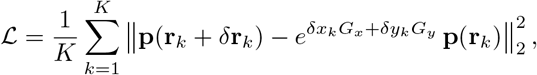

over a batch of *K* = 32 pairs per iteration, with start bins **r**_*k*_ drawn uniformly over the map and steps *δ***r**_*k*_ of up to two bins along each axis, reflected at the boundary. Short steps are kept because longer ones wrap the rotation angles and make optimization erratic. *B* was initialized uniformly in 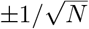 and the wavevectors from a Gaussian of standard deviation 2 in units of inverse arena half-width, and both were optimized with the Adam optimizer [55] for 20,000 iterations at learning rates of 10^−3^ for *B* and 10^−2^ for the wavevectors, with seed 0.

#### Reconstruction

Rate maps are regenerated by transporting the recorded population vector at the arena center to every bin, 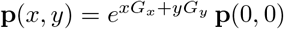, and compared with the data unit by unit through the Pearson correlation over bins. The median correlation is 0.76 for grid cells (10th to 90th percentile 0.59 to 0.83) and 0.77 for place cells (0.65 to 0.88), 0.82 for the units with a single field and 0.75 for those with several. Panel a of Figs. 9 and 10 shows every unit in order of correlation. The place-cell maps are noisier than the grid-cell maps, and the model reproduces the field and its surroundings while leaving out the finer structure of the data.

**Figure 9:**
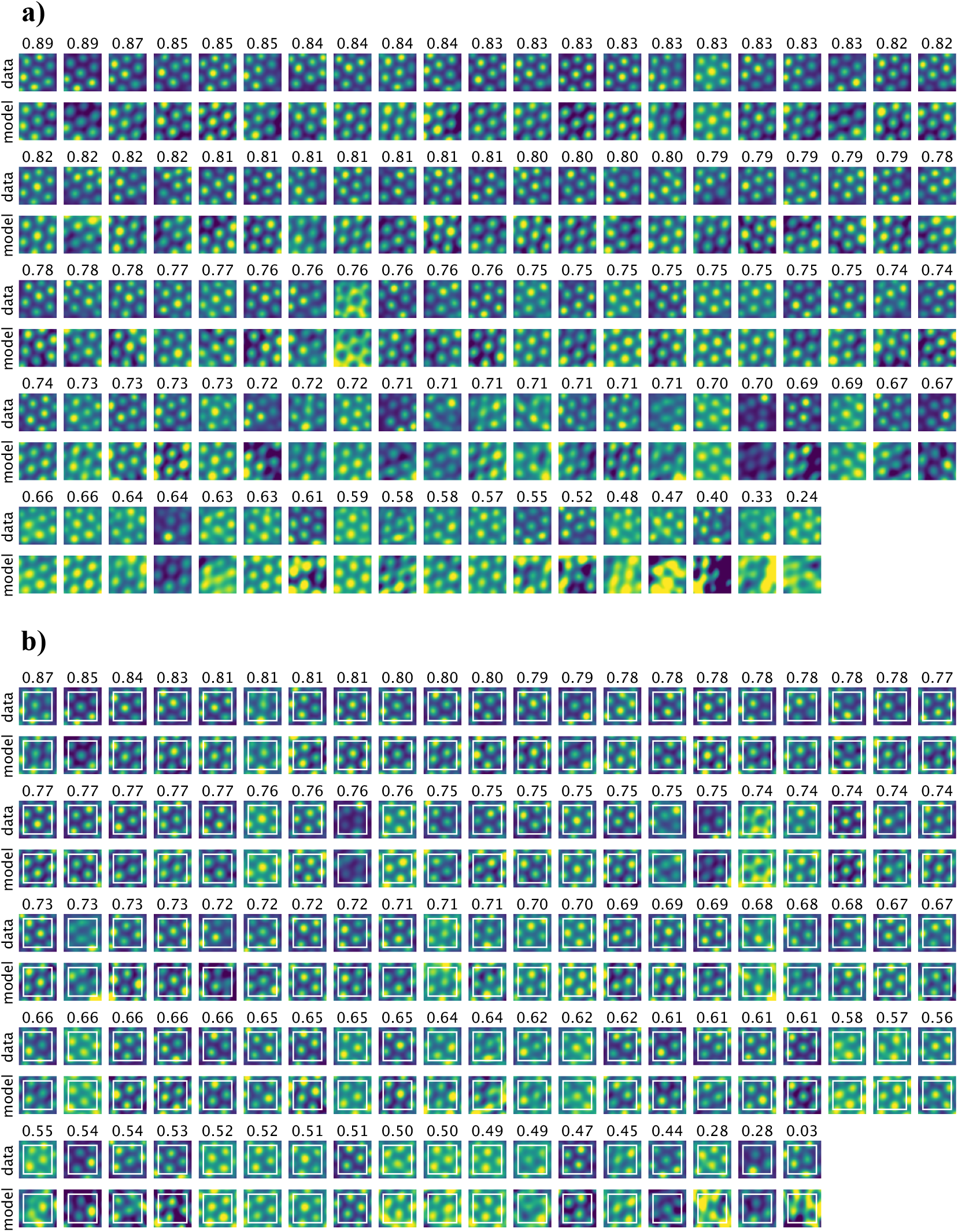
All 102 grid cells. **a)** Recorded rate maps [39] (data) and the maps regenerated by the fitted model (model), ordered by the correlation between the two, given above each pair. **b)** The model fitted to the central three quarters of the arena side (white square) and the full map generated from the center, ordered by the correlation between data and model over the part of the arena not trained on.

**Figure 10:**
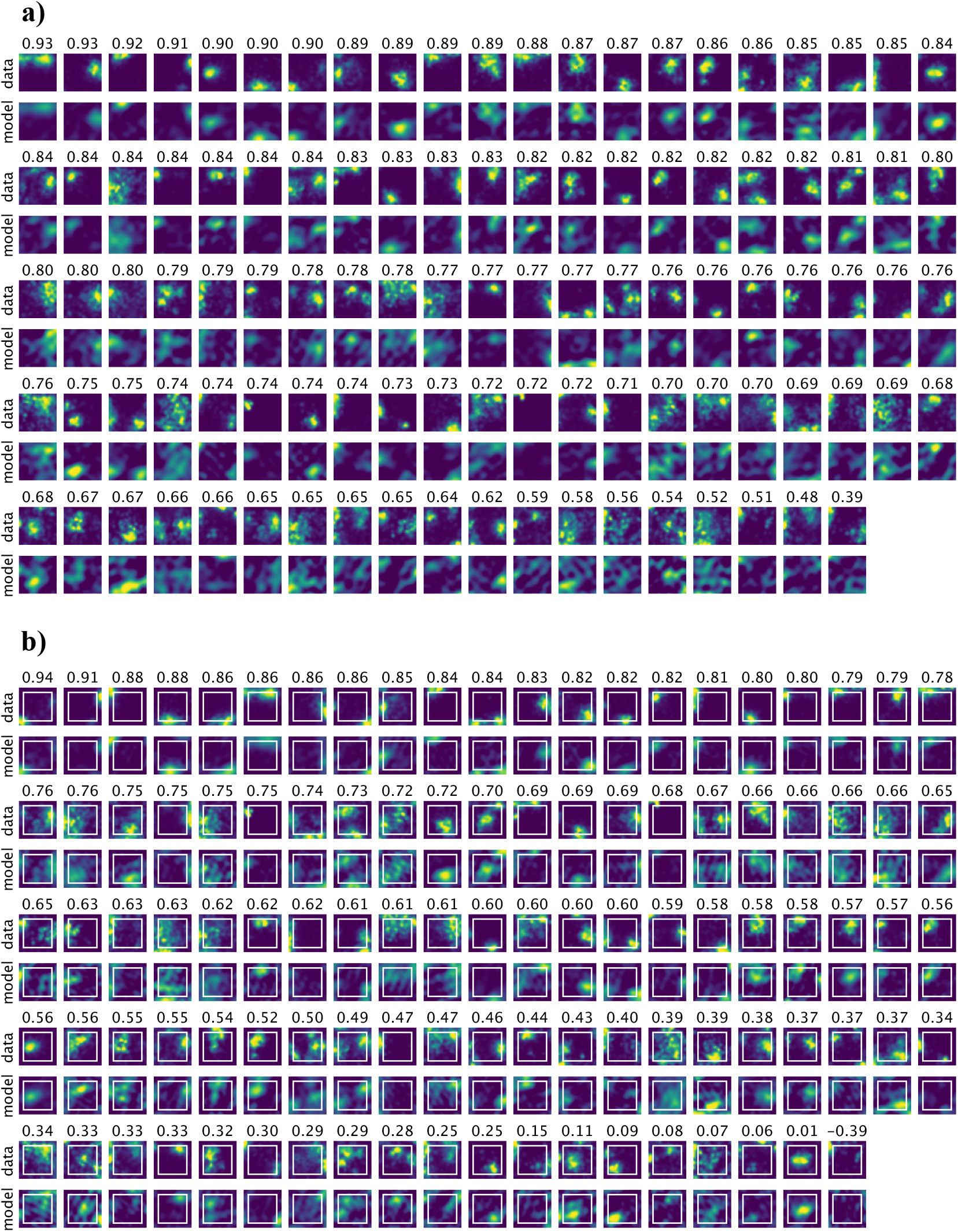
All 103 CA1 place cells. **a)** Recorded rate maps [40] (data) and the maps regenerated by the fitted model (model), ordered by the correlation between the two, given above each pair. **b)** The model fitted to the central three quarters of the arena side (white square) and the full map generated from the center, ordered by the correlation between data and model over the part of the arena not trained on.

#### Extrapolation with ground truth

To test extrapolation against recorded data, the model was fitted anew with the same protocol to a central square spanning three quarters of the arena side, bins 4 to 27 of 32 along each axis, so that the outer four bins on every side, 44% of the arena, were never used in training. The full map was then generated from the center and compared with the data over the untrained bins only. The median correlation over the untrained part is 0.71 for grid cells (0.51 to 0.80) and 0.61 for place cells (0.28 to 0.84), while the correlation inside the trained square is 0.92 and 0.81. The chance level is zero when the spatial bins of each recorded map are shuffled, and −0.03 and −0.02 when each unit’s prediction is scored against the data of a random other unit (200 draws each). Panel b of Figs. 9 and 10 shows every unit. The 49 of 103 place cells whose rate map peaks in the untrained part are predicted at a median correlation of 0.62, the same as the rest. Counting a recorded field as recovered when a model field, delineated by the same criterion relative to the unit’s recorded peak, covers at least half of its untrained bins, fields that extend into the trained square are continued outward in 221 of 229 cases for grid cells and 105 of 134 for place cells. Fields lying entirely in the untrained part are recovered for 21 of 29 grid fields but only 3 of 22 place fields, since the trained square determines the lattice but carries no information about where an isolated field lies.

#### Beyond the arena

Finally, the models fitted to the full arena were used to generate maps over an arena of twice the side length, centered on the recorded one, by the same transport from the center (Fig. 11). No data exist there, so these maps are predictions. For grid cells the hexagonal lattice continues across the boundary of the recording enclosure. For place cells all but one unit have fields lying entirely outside the recorded arena, four in the median by the field criterion above, with fields covering 31% of the area inside the enclosure and 30% outside, and outside peaks at 99% of the fitted peak rate in the median, though for a tenth of the units they exceed it by half or more. This follows from the construction rather than from the particular fit. The population vector keeps its norm under orthogonal transport, so the activity that forms a unit’s fields inside the arena cannot vanish outside it and reappears as further fields. The framework therefore predicts that place cells acquire further fields in a larger environment, the 47 units with a single field in the 1.5 m arena included, in line with the multiple fields reported for CA1 cells in large environments [41, 56]. The positions of the predicted fields are not constrained by the data, and recordings of the same cells in an expanded arena could be a direct test of this modeling approach.

**Figure 11:**
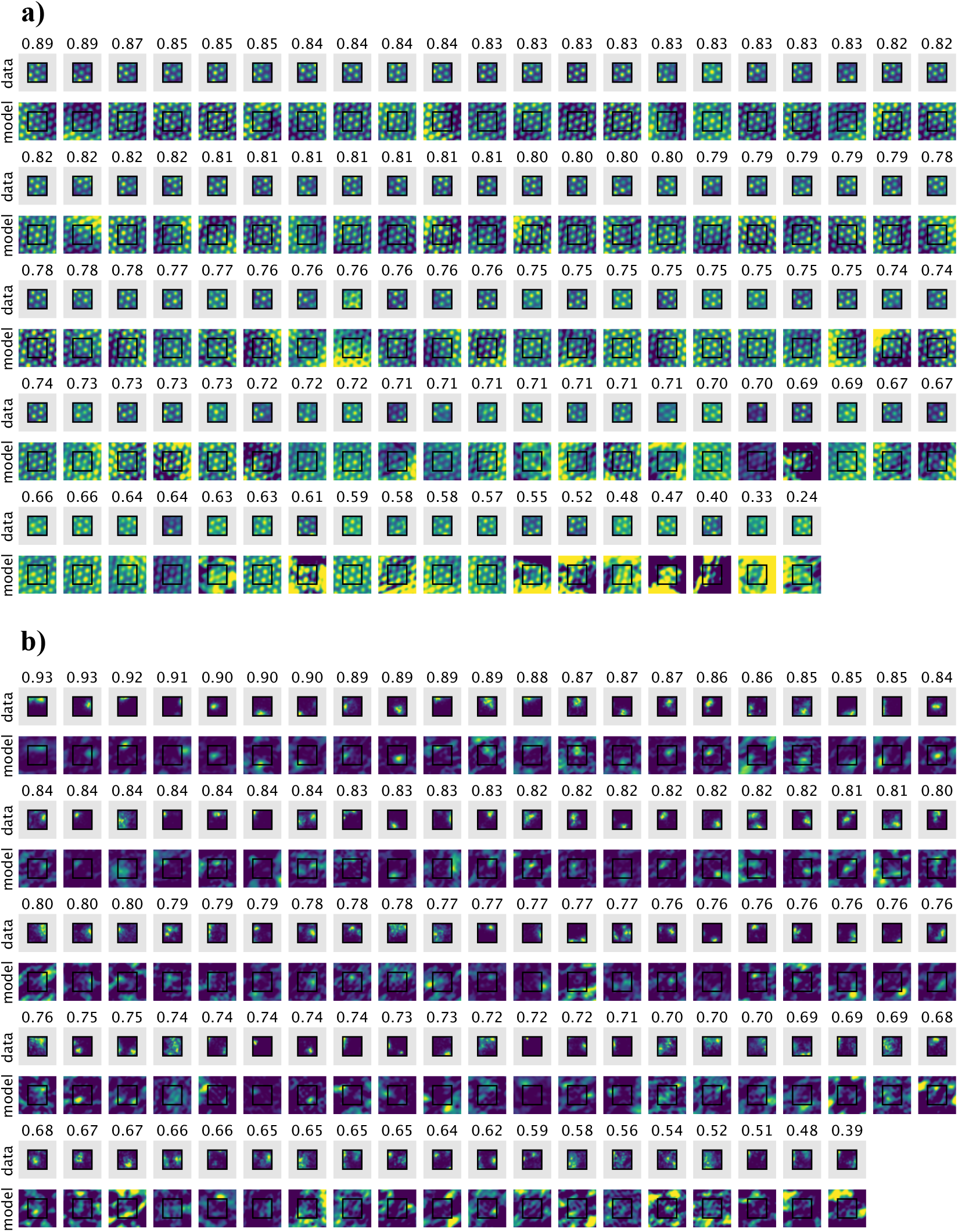
Generated maps over an arena of twice the side length. **a)** All 102 grid cells and **b)** all 103 place cells, ordered by the reconstruction correlation given above each pair. The recorded rate map (data) is shown in a grey frame at the position of the recording enclosure, outlined in black, above the map generated by the model fitted to the full arena (model). The model continues the grid lattice and places further fields of the place cells outside the enclosure.

#### Similarity structure

The fitted generators are skew-symmetric and commute, so the cosine similarity between the population vectors at two positions depends on their displacement alone (Eq. (3)), which the recordings need not satisfy. Figure 12 compares the two for the models fitted to the full arena. The recorded similarity averaged over all pairs of bins with the same displacement, up to half the arena side along each axis, is matched by the fitted similarity function at a correlation of 0.94 for grid cells and 0.98 for place cells (panels a, d). For grid cells both show the lattice of the module, and for place cells both fall off over about half the arena, with the fitted function somewhat elongated along one diagonal where the recorded average is only slightly so. Over individual pairs of bins the recorded similarity scatters around the fitted function (b, e), which explains 54% and 76% of its variance, against 65% and 81% for the displacement average itself.

**Figure 12:**
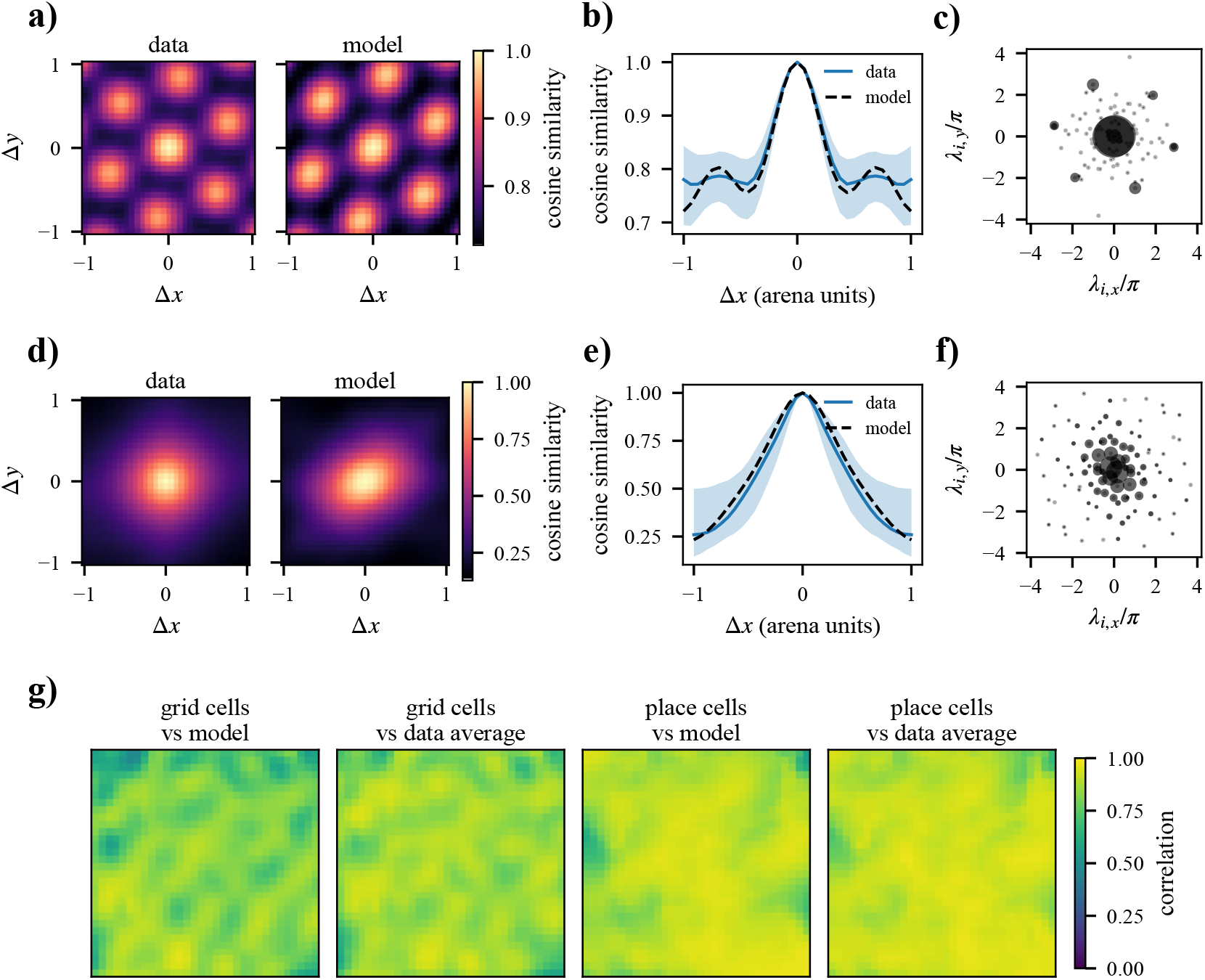
Similarity structure of the recordings and of the fitted models. **a)** Cosine similarity between the population vectors at two bins as a function of their displacement, for the recorded grid cells averaged over all pairs of bins with that displacement (data) and for the fitted model (model), on one color scale. **b)** The same similarity between bins displaced by (Δ*x*, 0), median and 10th to 90th percentile over all such pairs for the data and the single curve of the model. **c)** The fitted wavevectors (*λ*_*i,x*_, *λ*_*i,y*_) as grey dots at ±***λ***_*i*_, in cycles per arena side, with the area of each black disc proportional to a block’s share of the similarity function in Eq. (4). **d), e), f)** As a) to c) for the place cell data and fit. **g)** For every reference bin, the correlation between the recorded similarity map from that bin and the model’s (vs model), or the recorded similarity averaged over displacement (vs data average), over the bins within half an arena side.

Correlating the recorded similarity map from each reference bin with the model’s (g) gives a median of 0.82 for grid cells and 0.90 for place cells, against 0.86 and 0.92 for the displacement average, so the recorded similarity is close to translation invariant over most of the arena and departs from it for grid cells near the walls and corners. Together, these results suggest that the data are well described by a translation-invariant similarity structure.

The norm of the recorded population vector varies by 18% and 21% over the arena (standard deviation over mean), which the model cannot reproduce. Finally, Eq. (4) assigns each fitted wavevector a share of the similarity function, the weight 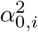 summed over its block (c, f). For grid cells 0.75 of the weight sits on a nearly constant component, as expected for non-negative rates that are not sparse, and 0.16 on four blocks at 2.7 to 2.9 cycles per arena side with orientations of 47^°^, 112^°^ and 170^°^, about 60^°^ apart, one hexagonal set as in Section 2.4. For place cells the weight is spread over 21 blocks of up to about 3 cycles per arena side with no preferred scale or orientation, similar to the construction in Appendix H. Note, however, that for place cells these similarities describe rate maps pooled over sessions rather than a simultaneously recorded population.

### L Binding and bundling as a basis for multiple-map reward-oriented navigation

While our primary focus has been on the geometric and computational properties of spatial representations, navigation involves more than localization. The algebraic structure of our spatial representations is deeply tied to hyperdimensional computing (HDC) [32], supporting robust goal-oriented navigation via bundling and binding.

HDC encodes information using distributed representations based on high-dimensional vectors, commonly termed hypervectors. A core property of high-dimensional spaces is the concentration of measure, which ensures that randomly sampled vectors become nearly orthogonal with high probability with increasing vector dimension. HDC systems exploit the statistical phenomenon via two primary operations. The first is bundling or superposition which aggregates vectors to form a composite representation that remains similar to the inputs. The second is binding, an operation that combines vectors to produce a result that is dissimilar and effectively orthogonal to its constituents.

#### L.1 Bundling and constructing reward maps

Assuming a similarity-preserving representation (such as the Gaussian similarity derived in Appendix I), we can implement a simple memory mechanism by bundling the population vectors of salient locations. Let 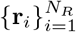 be a set of locations associated with a reward. A composite memory vector **p**_*R*_ is formed by summing the representations

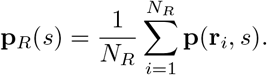

Querying the current location **p**(**r**, *s*) against the memory yields a similarity score *C*_*R*_(**r**, *s*) = **p**(**r**, *s*)^*T*^ **p**_*R*_(*s*) given by the equation

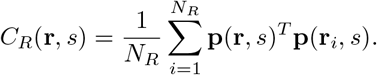

When we sample eigenvalues according to the process outlined in Appendix I, the pairwise similarity approximates a Gaussian, and the resulting score 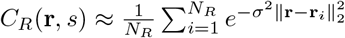 acts as a smooth reward density map resembling a mixture of Gaussians. The continuous landscape allows an agent to navigate to rewards by sampling local steps and following the gradient of the similarity surface using **r**_*t*+1_ ← **r**_*t*_ + *η*∇_**r**_*C*_*R*_.

#### L.2 Binding and context-dependent map retrieval

While bundling creates a single reward map, complex navigation requires storing distinct maps for distinct task demands within the identical physical environment. The standard HDC binding operation orthogonalizes vectors. In our formalism, context-dependent remapping functions as an intrinsic binding operation.

Recall that the context generator *G*_*s*_ produces orthogonal transformations. If the remapping is sufficiently strong, implying dissimilar contexts, the representations become nearly orthogonal. Context signals orthogonally bind spatial representations across distinct task demands within the identical physical environment. We can form a general context-dependent memory vector **p**_*R,S*_ that bundles spatial locations across distinct task states *s*_1_ and *s*_2_ evaluating to

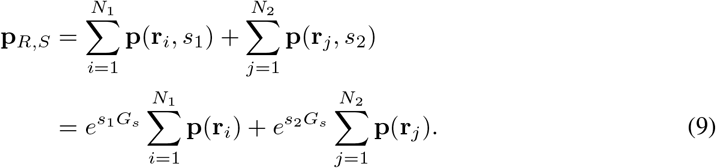

When we query the composite memory with a current state in the foraging context *s*_1_, the orthogonality of the remapping mathematically filters out the interference from the homing context *s*_2_. The query operation reduces to

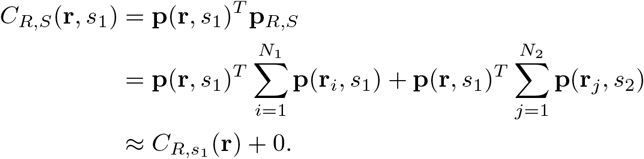

Here the cross-context term evaluates to zero because the relative context shift Δ*s* = *s*_2_ − *s*_1_ decorrelates the vectors. The explicit cancellation allows multiple reward maps to be superimposed within a single neural population in a robust manner facilitating context-specific retrieval. Consequently an agent can transition from foraging to homing behavior within the same arena solely by shifting the context signal and querying the composite memory.

### M Pairwise phase relations under a context shift

The spatial phase relations between grid cells recorded together are preserved under remapping [11], and relationships between pairs of grid cells with similar spacing are conserved even when the responses of single cells deform [45]. Here we derive the condition under which the exponential map has this property. We measure the relative spatial phase of two units by their spatial cross-correlogram, and ask when it is unchanged by a shift of the context signal *s*.

We assume a context generator *G*_*s*_ that is skew-symmetric and commutes with *G*_*x*_ and *G*_*y*_. The three generators then share the block-diagonal form of Appendix C, and the rotation angle of the *i*-th two-dimensional subspace becomes Ω_*i*_ = **k**_*i*_ · **r** + *λ*_*i,s*_*s*, with position **r** = (*x, y*) and wavevector **k**_*i*_ = (*λ*_*i,x*_, *λ*_*i,y*_). Since **p** = *R*^*T*^ ***α***, the activity of unit *n* is a sum of one plane wave per subspace,

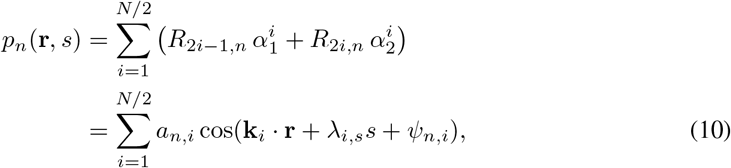

with amplitude 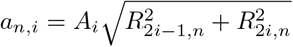 and phase offset *ψ*_*n,i*_ = *ω*_*i*_ − *β*_*n,i*_, where *β*_*n,i*_ is the polar angle of (*R*_2*i*_−_1,*n*_, *R*_2*i,n*_). The mixing matrix *R* and the origin state **p**_0_ enter only through *a*_*n,i*_ and *ψ*_*n,i*_. A context shift adds the phase *λ*_*i,s*_*s* to the *i*-th plane wave, and this phase is the same for every unit.

We define the spatial cross-correlogram of units *n* and *m* in context *s* as

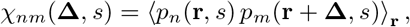

where the average runs over a region that is large compared to the grid period. We take the wavevectors to be nonzero and to lie in a half-plane, as for the rings of Section 2.4. Inserting Eq. (10) and using 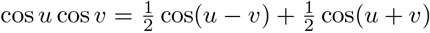 gives terms that oscillate in **r** with wavevectors **k**_*i*_ − **k**_*j*_ and **k**_*i*_ + **k**_*j*_. The spatial average of such a term vanishes unless its wavevector is zero, which happens only for **k**_*i*_ = **k**_*j*_. Separating the terms with *i* = *j* from those where two distinct subspaces share a wavevector, we obtain

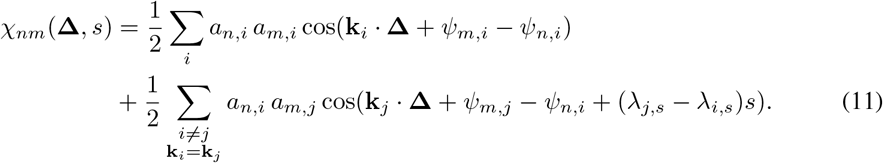

In the first sum the context phases cancel, because both units acquire the same phase *λ*_*i,s*_*s* in the *i*-th subspace. This sum is independent of *s* for any mixing matrix *R*, any origin state **p**_0_ and any context eigenvalues. Each term of the second sum oscillates in *s* with frequency *λ*_*j,s*_ − *λ*_*i,s*_, and for a dense mixing matrix its amplitude *a*_*n,i*_*a*_*m,j*_ is in general nonzero. The cross-correlogram of every pair of units is therefore independent of the context when

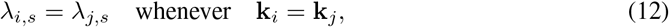

so that subspaces that share a wavevector also share a context eigenvalue. The variances *χ*_*nn*_(**0**, *s*) have the same form, and the normalized cross-correlogram is then unchanged as well. If every wavevector belongs to a single subspace, the second sum is empty and Eq. (12) holds automatically.

Note that the fields of single units can still deform, that is, remap. Under Eq. (12) the subspaces of one wavevector rotate together, and a unit of a hexagonal module reduces to three plane waves with **k**_1_ − **k**_2_ + **k**_3_ = 0 and context eigenvalues *λ*_1,*s*_, *λ*_2,*s*_ and *λ*_3,*s*_. A rigid translation of the field by a displacement **d** shifts the phase of the *l*-th wave by **k**_*l*_ · **d**. A context shift is such a translation only if **k**_*l*_ · **d** = *λ*_*l,s*_*s* for all three waves, which requires

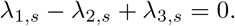

Otherwise the phase combination *ψ*_*n*,1_ − *ψ*_*n*,2_ + *ψ*_*n*,3_, which no translation can change, shifts by (*λ*_1,*s*_ − *λ*_2,*s*_ + *λ*_3,*s*_)*s* in every unit, and the fields deform. The lattice periodicity is unaffected, since the wavevectors do not depend on *s*. In both cases the pairwise cross-correlograms are unchanged.

The condition also constrains the similarity between contexts. At a fixed location, the similarity of a representation that satisfies Eq. (12) contains one cosine cos(*λ*_*l,s*_Δ*s*) per distinct wavevector. With the three wavevectors of a single hexagonal module, it does not decay as a Gaussian. Rigid pairwise relations and a Gaussian decay can hold together when the decay is carried by many distinct wavevectors. The example of Fig. 3 is of this kind. Its 50 hexagonal sets have distinct orientations, so every wavevector belongs to a single subspace, the Gaussian decay is carried by 150 context eigenvalues, and the pairwise cross-correlograms within each set are unchanged between contexts.

## Footnotes

1 Measured relative to a continuous positive baseline.

